# Pattern similarity analyses of frontoparietal task coding: Individual variation and genetic influences

**DOI:** 10.1101/642397

**Authors:** Joset A. Etzel, Ya’el Courtney, Caitlin E. Carey, Maria Z. Gehred, Arpana Agrawal, Todd S. Braver

## Abstract

Pattern similarity analyses are increasingly used to characterize coding properties of brain regions, but relatively few have focused on cognitive control processes in FrontoParietal regions. Here, we use the Human Connectome Project (HCP) N-back task fMRI dataset to examine individual differences and genetic influences on the coding of working memory load (0-back, 2-back) and perceptual category (Face, Place). Participants were grouped into 105 MZ (monozygotic) twin, 78 DZ (dizygotic) twin, 99 non-twin sibling, and 100 unrelated pairs. Activation pattern similarity was used to test the hypothesis that FrontoParietal regions would have higher similarity for same load conditions, while Visual regions would have higher similarity in same perceptual category conditions. Results confirmed this highly robust regional double dissociation in neural coding, which also predicted individual differences in behavioral performance. In pair-based analyses, anatomically-selective genetic relatedness effects were observed: relatedness predicted greater activation pattern similarity in FrontoParietal only for load coding, and in Visual only for perceptual coding. Further, in related pairs, the similarity of load coding in FrontoParietal regions was uniquely associated with behavioral performance. Together, these results highlight the power of task fMRI pattern similarity analyses for detecting key coding and heritability features of brain regions.

## Introduction

A current focus of recent cognitive neuroscience research has been to understand the functional specializations associated with brain networks, rather than focal brain regions. One network that has received a great deal of research attention is the frontoparietal network (FPN), based on a strong theoretical consensus that this network plays a critical role in higher cognitive functions such as working memory and executive control (Michael W Cole & Schneider, 2007; Duncan, 2010; Vincent, Kahn, Snyder, Raichle, & Buckner, 2008). Consequently, a key goal has been to determine whether FPN functionality in higher cognition can be understood in terms of the specific information that is being coded within this network. Yet progress in understanding the specific coding properties associated with the FPN has been especially challenging, for a number of reasons. One is related to an influential view of FPN, which postulates that this brain network is critical for higher cognition precisely because it has a highly flexible coding scheme, that is not fixed, but rather adapts to current task demands (Assem, Glasser, Van Essen, & Duncan, 2019; Duncan, 2001). A second reason is that the FPN is a brain network which seems to strongly reflect individual differences in cognitive functions and abilities. Indeed, a key characteristic of the cognitive functions attributed to the FPN, such as working memory and executive control, is that they are dominated by individual variation (Kane & Engle, 2002). Likewise, brain imaging studies have repeatedly shown that the FPN is the brain network most robustly associated with individual variation in higher cognitive functions (M. W Cole, Yarkoni, Repovs, Anticevic, & Braver, 2012; Michael W Cole et al., 2013; Jung & Haier, 2007; Kane & Engle, 2002). Consequently, it may be the case that understanding representational coding in the FPN needs to incorporate and account for such individual differences.

Thankfully, advances in cognitive neuroscience techniques have pointed to promising methods for investigating representational coding and individual differences in brain regions such as the FPN. Multivariate pattern analysis (MVPA) approaches may be particularly suitable for addressing such questions, given that they enable examination of information encoded in a distributed fashion, such as in large-scale brain networks. Prior work has used MVPA to demonstrate that information related to working memory and executive control, such as task rules, can be decoded from activation patterns within the FPN (Bode & Haynes, 2009; Michael W Cole, Etzel, Zacks, Schneider, & Braver, 2011; Etzel, Cole, Zacks, Kay, & Braver, 2016; Reverberi, Görgen, & Haynes, 2012; Waskom, Kumaran, Gordon, Rissman, & Wagner, 2014; Woolgar, Hampshire, Thompson, & Duncan, 2011; Woolgar, Thompson, Bor, & Duncan, 2011; Zhang, Kriegeskorte, Carlin, & Rowe, 2013). Moreover, we have recently shown that individual differences in FPN coding can predict variability in behavioral performance in executive control tasks (Etzel et al., 2016). A particular type of MVPA, pattern similarity analysis, also referred to as correlational MVPA (Hendriks, Daniels, Pegado, & de Beeck, 2017) or representational similarity analysis (Kriegeskorte, Goebel, & Bandettini, 2006; Nili et al., 2014), may be particularly appropriate for examination of individual differences in FPN coding, since it provides a direct measurement of the similarity of activation patterns, both between individuals and within individuals across tasks. Although pattern similarity approaches have been less frequently applied to research on the FPN and executive control, one recent study did demonstrate a tight coupling between idiosyncratic activation patterns within the lateral prefrontal cortex and specific attentional control strategies and task performance (Lee & Geng, 2017).

Another well-established experimental and methodological approach for exploiting individual differences in brain activation is the examination of twin or sibling similarity in task activation patterns, which suggest underlying genetic contributions to cognitive function. The primary logic underlying such studies is the assumption that if individual differences in brain activation patterns are genetically encoded, then they should be similar across related individuals, and highest in monozygotic (identical; MZ) twins, since they, on average, share 100% of their segregating loci. Indeed, the standard logic of the “twin design” is to directly compare the similarity of brain activation in MZ and dizygotic (fraternal; DZ) twins, since differences provide a direct estimate of the heritability of brain activation patterns, under certain assumptions (Polderman et al., 2015). A number of brain imaging studies have been conducted using twin designs, including some that have focused on working memory and executive control (Blokland et al., 2011, 2017; Koten et al., 2009; Matthews et al., 2007). This work has demonstrated that at least some proportion of the FPN activation variability that is generally presumed to be idiosyncratic is in fact heritable, and thus a meaningful component of individual differences. An exemplar of this type of research is the work by Blokland and colleagues using a large-sample dataset from the Queensland Twin Imaging Study (Blokland et al., 2008, 2011, 2017). In a series of studies, these researchers used whole-brain genetic modeling techniques to demonstrate significant heritability effects (averaging 33% of variance) on brain activation patterns evoked during N-back task performance, which were primarily observed in FPN regions.

However, much of this previous work has focused on describing brain activity with univariate statistics: analyzing voxels individually (e.g., (Blokland et al., 2011)) or averaging across voxels within regions of interest (ROIs; e.g., (Blokland et al., 2008)). A limitation of this approach is that it is unable to detect patterns spanning multiple voxels. This is problematic because, as was alluded to above, the neural encoding of cognitive control-related representations is standardly thought to occur within such multivariate and distributed patterns of activity. Consequently, univariate approaches may miss some of the key dimensions of individual difference that may be present in distributed FPN activity patterns. A few neuroimaging studies have utilized multivariate approaches to test for heritability effects in twin designs (Pinel et al., 2015; Polk, Park, Smith, & Park, 2007). In this work, the key approach is to examine the relative similarity of activation patterns in MZ twins, relative to both DZ twins and unrelated individuals. In a first study of this type, Polk et al. (2007) showed that pattern similarity was reliably higher in members of MZ twin pairs within occipitotemporal regions when processing face and place stimuli. In a follow-up study by Pinel et al. (2015), this finding was confirmed within a face region, and was further found to differentiate from univariate approaches, which were not sensitive to a significant MZ similarity effect in this same region. This aspect of the Pinel et al. (2015) findings suggests that multivariate approaches may have potentially greater statistical and inferential power for detecting the heritability component of individual differences. Nevertheless, to our knowledge, no prior studies have used multivariate approaches to estimate heritability and other familial effects and individual differences within FPN regions, through twin-based designs.

Another related question refers to the functional-anatomic specificity of twin-based multivariate pattern similarity effects. Although the prior studies focusing on occipitotemporal regions have shown the potential power of multivariate approaches for detecting twin similarity effects in perceptual coding, other brain regions were not tested, so the anatomic specificity of genetically-based pattern similarity for the same perceptual categories (i.e., faces and places) was not evaluated. In other words, although it is commonly assumed that the multivariate pattern similarity effects occur preferentially in the brain regions thought to be involved in perceptual coding, i.e., occipitotemporal regions, this assumption has not been tested directly, by comparison against other brain regions that would not be thought to mediate such coding (e.g., FPN). Conversely, in studies examining FPN twin pattern similarity effects, it would be important to test whether such effects preferentially occur in conjunction with task conditions that do involve FPN coding, such as working memory and executive control.

A final limitation of the prior work is that it has not provided strong tests of whether activation pattern similarity effects are functionally relevant, for example, by linking them to behavioral performance. In particular, if activation pattern similarity reflects the fidelity or quality of representational coding, then individual differences in activation pattern similarity should predict variation in behavioral performance. Moreover, twin pairs showing higher activation pattern similarity (i.e., with each other) should also be more likely to have better task performance. This prediction derives from the assumption that activation pattern dissimilarity reflects, in some part, noisiness in representational coding – since by definition noisy representations should be uncorrelated across individuals, whereas optimal representations should be more similar among related pairs.

In the current study, we systematically tested each of these predictions, addressing limitations in the prior work. Specifically, we exploited the large size and family-based structure of the Human Connectome Project (HCP, (Van Essen et al., 2012)) dataset to investigate individual differences and twin similarity in neural representational coding. We focused on two brain networks for which the prior literature suggests the most clear-cut predictions of functional specialization and dissociability: the FPN and a contrasting Visual occipitotemporal network. We took advantage of the richness of the N-back task fMRI within the HCP to examine representational coding of working memory (WM) load and perceptual category. In the HCP, participants performed the N-back under high (2-back) and low load (0-back) conditions with multiple categories of image stimuli, including faces and places. Neural coding related to working memory load should result in higher activation pattern similarity across conditions that have the same WM load but different perceptual categories (e.g., 2-back Face and 2-back Place) compared to conditions that have different WM loads (e.g., 2-back Place and 0-back Place). In the same manner, similarity-based approaches can be used to test for anatomical specificity, since WM load-based coding should be present in FPN but not visual occipitotemporal brain regions; conversely, perceptual category coding should be present in visual occipitotemporal brain regions but not FPN. As a first step of analysis, we tested for double dissociability of neural coding to validate the pattern similarity analysis approach as a means of addressing individual differences questions within the FPN.

After establishing such effects, the second set of analyses tested the hypothesis that pattern similarity in MZ twins would be greater than in relatives that are less genetically similar (i.e., DZ twins and non-twin sibling pairs) and further, within unrelated pairs who also do not share other familial influences (e.g., shared environment). Even more critically, we provide a novel test of the anatomical specificity present in this pattern as well, such that FPN would exhibit high pattern similarity among twins selectively for load-based coding, whereas in visual occipitotemporal regions high twin-based pattern similarity would be selectively present for perceptual category coding. Finally, we examined whether variability in pattern similarity was functionally meaningful, in the sense of predicting better N-back task performance, and moreover whether twin pair-based variation could uniquely predict performance over and above individual variation. To preview, our findings strongly confirmed each of these predictions, providing a clear base of support for the idea that FPN activation patterns reflect functionally specific coding of WM-related information, while also incorporating a substantial degree of both individual and genetically-related variation.

## Materials & Methods

### Dataset and Participants

This work used functional images, demographic information, and behavioral performance measures from the Human Connectome Project (HCP, http://humanconnectome.org/) (Van Essen et al., 2012). We included a total of 764 participants (382 pairs) from the 1200 subjects release, selected to form four groups of paired people: monozygotic twins (MZ; 105 twin pairs), dizygotic (DZ, 78 twin pairs), non-twin siblings (SIB, 99 pairs); and unrelated people (UNR, 100 pairs). Only twins with genomically-verified zygosity (as of March 2017) were included, and twin status was assigned based on this information, rather than self-report. Only same-sex pairs were included in each group. MZ and DZ twin pairs are necessarily of the same age (though sometimes scanned several months apart); pairs of non-twin siblings and unrelated people were selected to be within 3 years of age at the time of scanning. Only full siblings (same mother and father) were included in SIB; unrelated people did not share either a mother or a father by their self-report. No person was included in more than one group (i.e., a person would not be paired with their twin in the MZ group and someone else in the UNR group).

All analyses were performed using R version 3.1.3 (R Development Core Team, 2015), with WRS (Wilcox, 2017) and DescTools package functions for robust statistical tests. Trimmed (at .1) means and standard errors are reported unless otherwise specified. Code and input data for replicating the figures and analyses is in the Supplemental and available at the Open Science Foundation, https://osf.io/p6msu/.

### Task and Data Processing

We used functional images from the HCP working memory task fMRI dataset, which is a blocked version of the N-back task (Barch et al., 2013). Briefly, task stimuli consisted of visual images (faces, places, tools, and body parts), with each block composed of a single category of images, performed with either 0-back load (judge whether the currently presented image matches the target image shown at the beginning of the block) or 2-back load (judge whether the currently presented image matches the one shown two trials back). The task was presented in two imaging runs, each of which had eight task blocks, one block for each combination of load and stimulus category (Barch et al., 2013). Our analyses began with the parameter estimate images (second-level FSL COPEs (Smith et al., 2004)) included in the HCP 1200 subjects release. Briefly, these COPEs are from GLMs performed after the images went through the HCP Minimal Preprocessing Pipelines, which included (among other steps) MNI atlas transformation, surface projection, and surface parcel constrained 2 mm FWHM smoothing (Glasser et al., 2013).

Here, we focus on two distinct cognitive dimensions: perceptual category processing and working memory load processing. In the HCP working memory task, perceptual category processing varied with which type of image was used in a block, while working memory load processing varied with whether the block was 0-back or 2-back (higher load for 2-back). To focus on these processes in a balanced 2×2 design we included only face and place blocks in our analysis, with four parameter estimate images of interest for each person: 0-back Face, 2-back Face, 0-back Place, and 2-back Place. N-back load, face, and place image processing have been extensively studied, allowing clear-cut *a priori* predictions regarding the brain regions likely to be relevant. Specifically, we assumed that perceptual coding of perceptual category would occur within the ventral occipitotemporal visual network (Haxby et al., 2001) with fairly consistent patterns across individuals; conversely, coding of cognitive task goals and working memory load were assumed to occur within the frontoparietal control network (M. W Cole et al., 2012; Michael W Cole & Schneider, 2007; Owen, McMillan, Laird, & Bullmore, 2005; Vincent et al., 2008), and also be more idiosyncratic (highly impacted by individual differences).

As we could make strong expectations of which brain regions would be relevant for these cognitive processing components (the FPN and visual occipitotemporal networks), and since the goal of the current investigations was to explore signal strength and heritability patterns, rather than to isolate more anatomically localized brain regions (the question of anatomic specificity is an interesting one that we address briefly below, but a thorough investigation is beyond the scope of the current work), we employed a network-based analytic strategy. Specifically, we used the Gordon cortical parcellation scheme (Gordon et al., 2016) to anatomically identify the key regions of interest (ROIs) within the FPN and visual occipitotemporal networks. Following the terminology of this scheme, we hereafter refer to the specific sets of vertices that define these ROIs as FrontoParietal and Visual “communities”, rather than networks, reserving the term FPN to refer to the more generalized (i.e., not tied to a specific parcellation scheme) definition of this functional brain network.

The Gordon parcellation was released aligned to the HCP preprocessed images, making it convenient, while also unbiased with respect to our primary analyses. The two communities include brain regions that are generally considered relevant for working memory load and perceptual category processing (Figure 1, lower left). All vertices falling within each of these communities were included in the analyses; no further feature selection was performed. This resulted in, for FrontoParietal, 831 vertices in the left hemisphere and 1418 in the right hemisphere; for Visual, 3084 vertices on the left and 3689 on the right. Analyses were performed in each hemisphere separately, and then averaged over hemisphere.

**Figure 1.**
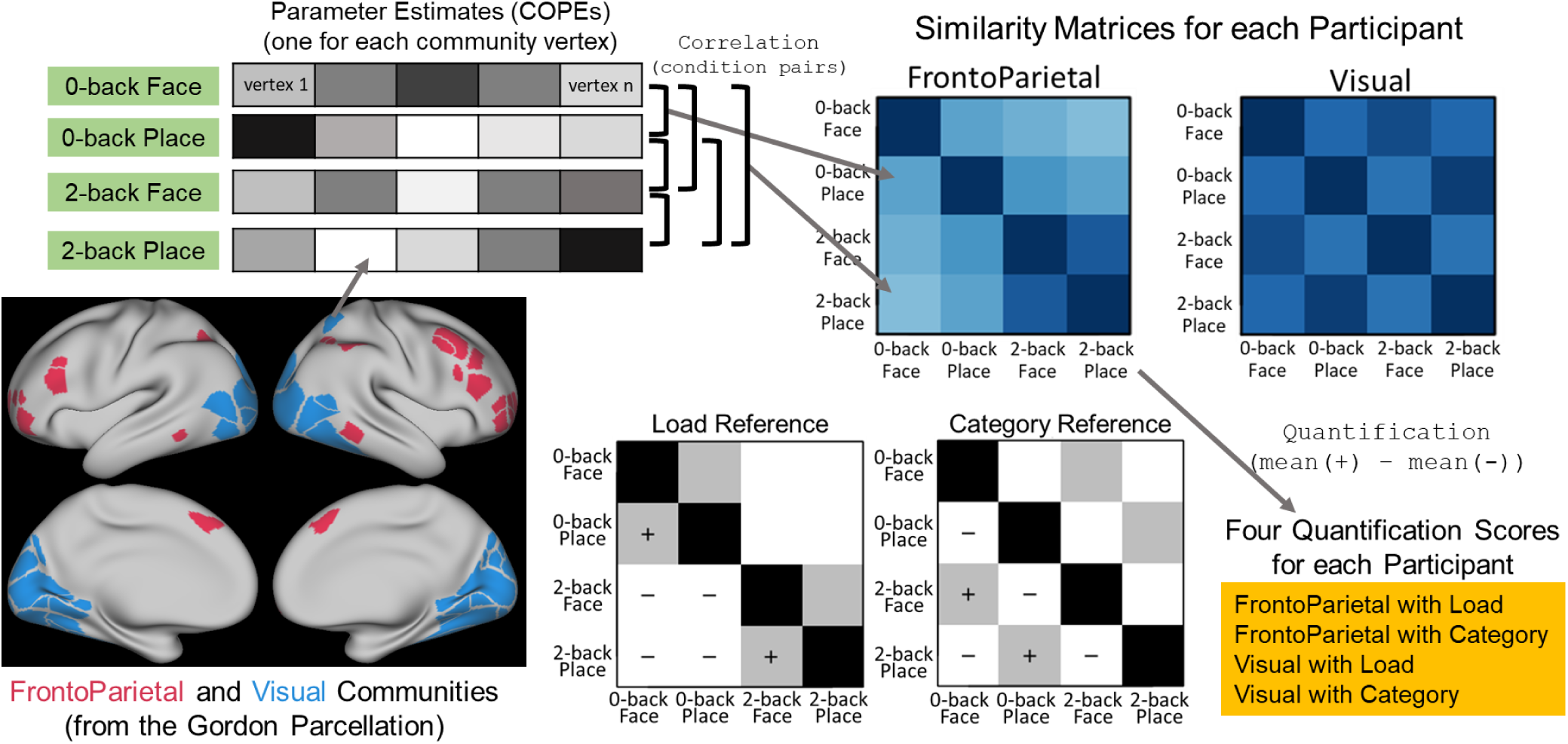
Illustration of the method of quantifying similarity within each individual participant. Starting at lower left and moving clockwise, we analyzed vertices within the Gordon FrontoParietal and Visual communities (Gordon et al., 2016). The values were extracted for each vertex from the parameter estimate images (COPEs, as released by the HCP), one vector for each of the four conditions of interest. The Pearson correlation between all pairs of these vectors was calculated, and arranged in the form of a similarity matrix; upper right. Two matrices, one for each community, were made for each participant. Finally, the Load and Category information in each matrix was quantified by subtracting the average of the cells marked with – from the average of the cells marked with + in the Reference Matrices, resulting in four scores for each participant.

### Load and Category Index Scores for Pattern Similarity Quantification

Pattern similarity approaches provide a means of quantifying the relative degree of similarity between activation patterns exhibited in different conditions, within brain regions of interest. For this study, our hypotheses concerned the relative similarity of vertex-level activity patterns between different task conditions that shared either the same perceptual Category (e.g., 0-back Face and 2-back Face) or the same working memory Load (e.g., 2-back Place and 2-back Face). Specifically, we expected that similarity related to Category would be higher in Visual than FrontoParietal, whereas similarity related to Load would be higher in FrontoParietal than Visual. We measured pattern similarity between conditions using the Pearson correlation statistic, following a common approach used in prior analyses (Haxby et al., 2001; Polk et al., 2007). Note that Pearson correlation is insensitive to additive and proportional translations: it ignores differences in the across-vertices mean value of each example, but will detect similarly-shaped vectors (e.g., higher values in vertex 2 than vertex 1) (Romesburg, 2004). Accordingly, no transformations (e.g., normalization) were made to the parameter estimate images: correlation was calculated using the HCP-released COPE value for each ROI vertex.

We conducted two sets of pattern similarity analyses, the first within individual participants (Figure 1), and the second within paired participants (e.g., a pair of MZ co-twins, Figure 2). In both types of analyses, the correlations can be arranged into matrices, as is the standard approach in Representational Similarity Analysis (RSA, (Kriegeskorte, Mur, & Bandettini, 2008; Nili et al., 2014)). Given the four conditions, six correlations are possible within each individual (0-back Face with 2-back Face, 0-back Face with 0-back Place, etc.), forming symmetric similarity matrices, while sixteen unique correlations are possible within each participant pair (0-back Face in one person with 2-back Face of their co-twin, etc.). The appearance of the similarity matrices themselves can be useful, as can be defining a statistic to describe the degree to which particular information coding schemes are reflected in each matrix. Reference Matrices illustrate the expected appearance for each particular type of information coding; the Reference Matrices for Load and Category coding for individuals are shown in Figure 1 and for paired participants in Figure 2. There are multiple approaches for calculating how well each observed RSA matrix matches a reference; here we computed a difference score by subtracting the mean of matrix cells specifying the task conditions that were predicted to be less correlated (marked with – in Figures 1 and 2) from the mean of cells for conditions that were predicted to be more correlated (marked +). This difference-based quantification method is sometimes described as applying a contrast, with reference matrix cells weighted to sum to zero (e.g., (Oedekoven, Keidel, Berens, & Bird, 2017)).

**Figure 2.**
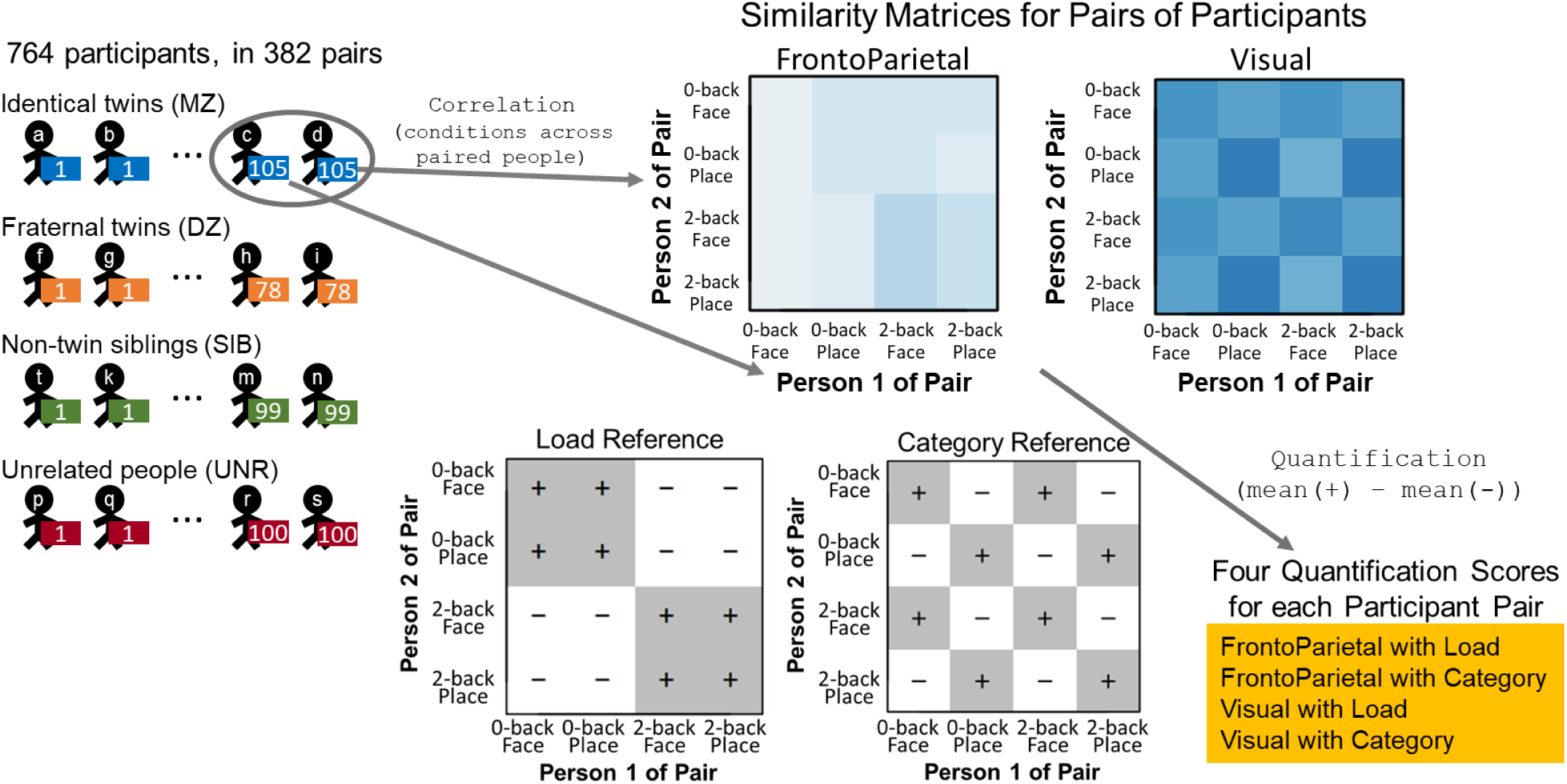
Illustration of the method of quantifying similarity of paired participants. The imaging data (parameter estimates) for each participant was extracted for each of the four conditions and two communities, as in Figure 1. The similarity matrix for each participant pair was constructed as Pearson correlations of all possible condition combinations between the two participants (e.g., 2-back Place of Person 2 correlated with 2-back Face of Person 1). Note that, unlike the similarity matrices for each individual participant, these matrices are not symmetric, and the diagonal is not 1. Finally, the Load and Category information in each matrix was quantified by subtracting the average of the cells marked with – from the average of the cells marked with + in the Reference Matrices, resulting in four scores for each pair of participants.

It is clear how to construct the Category reference matrices for both the individual and pairwise analyses: images of the same category should be more similar than images from different categories, with no expectation that the two image categories (Face and Place) would have different activation strength in these large communities. This equivalence does not hold for Load, however: within FrontoParietal the mean level of activation (i.e., a univariate statistic) generally increases as working memory load increases (although not a focus of the current work, we briefly explore these types of univariate activation effects in S5.2). Further, the HCP used a 0-back manipulation, which likely differs from the 2-back in more aspects than working memory load alone (i.e., 0-back and 2-back are likely to differ more than 2-back and 4-back would differ). It thus seems reasonable that we should only expect the similarity of two 2-back load conditions to be greater than the similarity of two conditions that differ in load. Restricting the Load quantification to 2-back conditions is not a perfect solution, however, because it unbalances the reference matrices, i.e., some cells are omitted from the Load quantification that are included in the Category quantification. Given this uncertainty regarding the best way to quantify Load, we settled on a conservative approach, including both 0-back and 2-back trials in the main analyses (as shown in Figures 1 and 2), but also conducting pairwise analyses including only 2-back trials for comparison. Thankfully, as described under Results, the primary findings were the same with both analysis, but did show evidence of greater sensitivity when only including 2-back trials.

## Results

### Behavioral Performance and Heritability

We first report behavioral task performance to validate expected patterns, both across the entire group, and in terms of heritability effects. N-back working memory performance was quantified in terms of d’ (Hautus, 1995; Pallier, 2002), proportion correct, and median reaction time (ms, calculated from correct trials only). First, we compared performance across the four participant groups (MZ, DZ, SIB, UNR), collapsing across condition (Table 1). The groups were not predicted to differ in performance, and in fact this was primarily the case. Groups did not differ in d’ (p=.08) or RT (p=.1), although DZ twins did show evidence of slightly better performance (p=.02, measured with proportion correct and t1way, a robust ANOVA (Wilcox, 2017); S1.1a). Next, we verified the presence of a significant load effect (poorer performance on 2-back relative to 0-back; p<.001 for all three measures). Again, there was no significant interaction with subject group or perceptual category (robust ANOVA; S1.1a). There was, however, a significant main effect of category (p=.038 for proportion correct, p<.001 for d’ and RT), such that responses were faster and more accurate for Face trials (relative to Place).

**Table 1.** Mean {standard error} of behavioral measures, all subjects combined, calculated from trials with the indicated conditions. Distributions and subject group separated results shown in S1.1.

|  | Face, Place<br>0-back, 2-back | Face, Place<br>0-back | Face, Place<br>2-back | Face<br>0-back, 2-back | Place<br>0-back, 2-back |
| --- | --- | --- | --- | --- | --- |
| Proportion Correct | .92 {.003} | .954 {.003} | .896 {.003} | .926 {.003} | .918 {.003} |
| d' | 2.6 {.04} | 3.01 {.04} | 2.23 {.04} | 2.62 {.04} | 2.44 {.04} |
| Median RT | 799.8 {4.8} | 700 {4.5} | 921.4 {5.8} | 787.7 {5.1} | 810.3 {5.1} |

Pairs were expected to show higher similarity as genetic similarity increased; thus, the three groups of related individuals (MZ, DZ, SIB) were predicted to have more similar performance than the unrelated pairs (UNR; see S5.1 for a control analysis in which UNR pairs were matched to have similar performance). Likewise, if genetic factors make a strong contribution to cognitive task performance, the MZ pairs would be predicted to show the strongest within-trait correlations. As shown in Table 2, we did find that all three related groups showed stronger similarity than the unrelated pairs for proportion correct and d’. The same trend was present for RT, but was only significant when comparing MZ twins to unrelated pairs. In all N-back performance measures, similarity was numerically highest for MZ twin pairs, but was not significantly different from the DZ or SIB pairs.

**Table 2.** Pearson correlation between the paired people in each subject group on the behavioral performance measures. p-values (in parentheses) were calculated with hc4wtest, a robust regression test for R^2^ different than zero (Wilcox, 2017), and uncorrected for multiple comparisons. Scatterplots and regression lines are in S1.2. Significantly different pairwise correlations within each row (i.e., on each measure) are indicated by shared superscripts, with p < .0083 (Bonferroni correction of .05 for 6 comparisons) as the significance threshold. All pairwise comparison p-values are listed in S1.2, and were calculated by twohc4cor (Wilcox, 2017). The astute reader will note negative correlations among the UNR pairs, which unexpectedly reached statistical significance for some of the measures. We believe the observed negative correlations reflect a sampling anomaly, as a larger set of unrelated pairings was quite close to the expected zero correlation (S1.5). Regardless, this does not seriously influence our key analyses or interpretations, which center on neural pattern similarity relationships among related pairs.

|  | MZ | DZ | SIB | UNR |
| --- | --- | --- | --- | --- |
| Proportion Correct | .44 ($p < .001$)<sup>a</sup> | .14 ($p = .13$)<sup>b</sup> | .36 ($p < .001$)<sup>c</sup> | -.32 ($p = .004$)<sup>a,b,c</sup> |
| d' | .43 ($p < .001$)<sup>a</sup> | .25 ($p = .03$)<sup>b</sup> | .32 ($p < .001$)<sup>c</sup> | -.38 ($p = .002$)<sup>a,b,c</sup> |
| Median RT | .34 ($p < .001$)<sup>a</sup> | .17 ($p = .062$) | .11 ($p = .388$) | -.07 ($p = .4$)<sup>a</sup> |

A parallel way to reveal the same point is through classic heritability modeling, which enables estimates of the proportion of variance that is genetic in origin. Using the classical twin model, as implemented with ACE structure (where A refers to additive genetic, C to common familial environment, and E to individual-specific environment; the correlations did not suggest the role of non-additive genetic factors)(Evans, Gillespie, & Martin, 2002), we estimated these parameters for the N-back behavioral measures. The best-fitting model by Akaike’s Information Criterion (AIC) was one in which A and E significantly contributed to variance in these measures, with heritability estimates in expected ranges (.36-.44) and statistically significant when estimates of common environment were constrained to zero, without any deterioration of fit (S1.6). These estimates are similar to what has been observed in prior heritability analyses of the N-back task (Blokland et al., 2008). We next examined the brain activity data to determine whether estimation of genetic factors could be detected with similar, if not higher, sensitivity and specificity than the associated behavioral measures.

### Anatomical Specificity of Activation Similarity Patterns: Analyses in individuals

The first analyses were conducted to establish sufficient power to detect heritability and validate that pattern similarity analysis methods were sufficiently sensitive to demonstrate the expected functional and anatomic specificity. We first considered which conditions should have more similar activation patterns for the two types of information coding: if working memory load is coded in a brain network, then conditions sharing the same load should exhibit similar activation patterns (**0-back** Face and **0-back** Place; **2-back** Face and **2-back** Place), while if perceptual category is coded, conditions sharing the same category should be more similar (0-back **Face** and 2-back **Face**; 0-back **Place** and 2-back **Place**). We expected that the FrontoParietal community would show more evidence of load-related than category-related similarity, with the reverse profile in the Visual community. Following the conventions of pattern similarity analysis, these load- and category-related similarity predictions are shown as Reference Matrices in Figure 1. Next, correlations among the six pairwise combinations of the four parameter estimate images were calculated for each community within each person, examples of which are in Figure 3 and S2.1. Visual inspection of individuals’ matrices suggest a clear difference between the communities: the FrontoParietal matrices tend to resemble the Load reference, while the Visual tend to resemble the Category reference. This impression was evaluated numerically by quantifying the Load and Category information in each individual’s Visual and FrontoParietal matrices, calculating differences according to the reference matrices (Figure 1), which provides four scores for each participant: FrontoParietal Load, FrontoParietal Category, Visual Load, and Visual Category.

**Figure 3.**
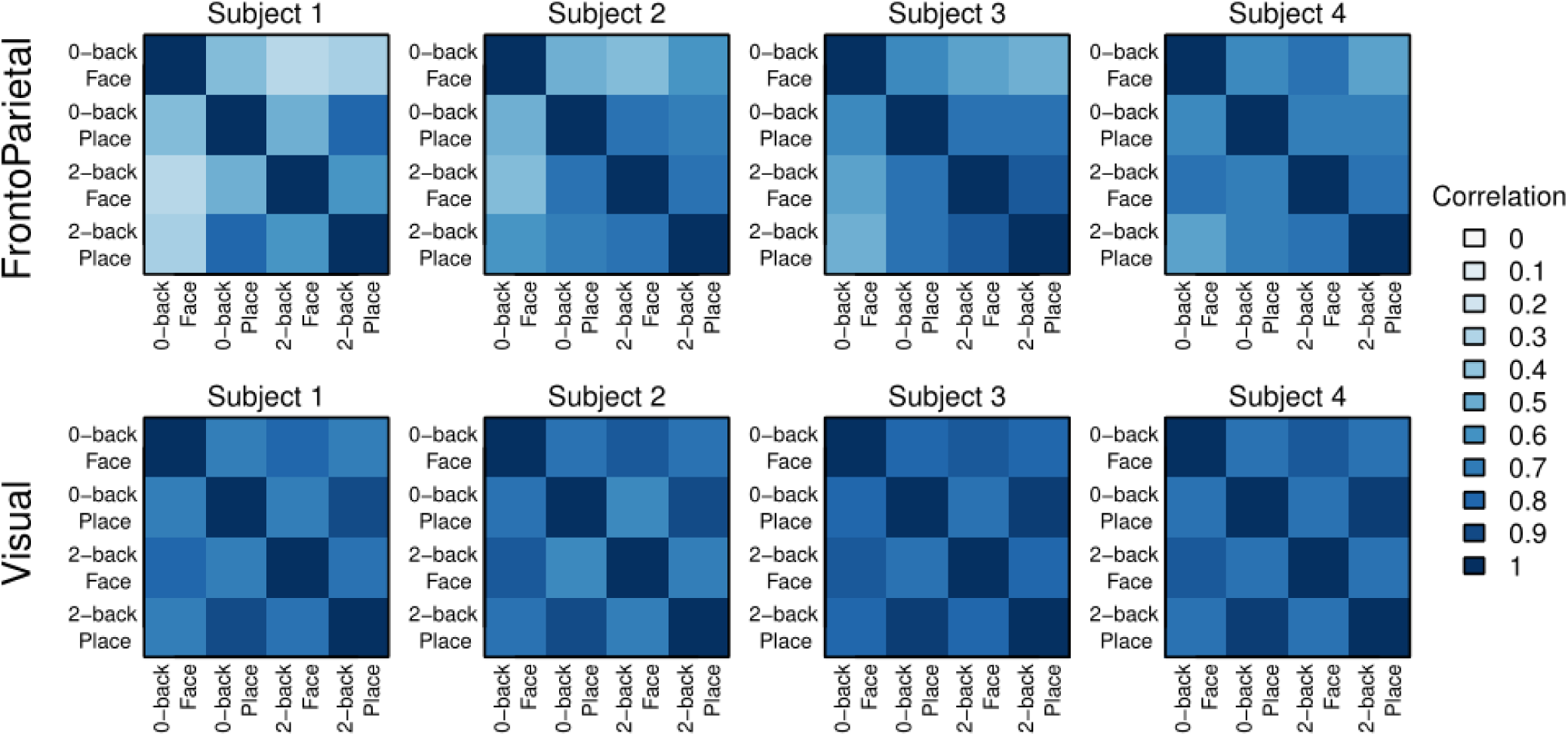
Similarity matrices for the two communities in four representative individuals. Additional examples are in Results Supplement 2, section S2.1, and group averages shown in S2.2. Note that the FrontoParietal matrices tend to resemble the Load Reference (Figure 1), while the Visual matrices tend to resemble the Category Reference, a tendency confirmed by the distribution of quantification scores (Figure 4).

The distribution of the four scores (Figure 4, S2.3) provides a clear indication of anatomical and functional specificity: in the FrontoParietal community, the quantification scores were significantly higher for Load than Category (p<.001), whereas in the Visual community the reverse pattern was present (Category > Load, p<.001). These patterns reflect a highly robust double dissociation (Community by Information Coding interaction, p<.001), supporting the notion of specificity in FrontoParietal as well as Visual (all pairwise contrasts also significant). Moreover, the prior analyses included all participants, but the double dissociation is highly significant within each separate participant group (MZ, DZ, SIB, UNR) as well (S2.3). In an exploratory follow-up analysis we tested for the same dissociation, but across the whole brain within each individual parcel, rather than only the two communities (S2.6). The parcel-level results were quite consistent with the community-level results, showing that Category > Load effects were primarily observed in Visual parcels, while Load > Category effects were most robust in FrontoParietal, DorsalAttention, and Default Mode parcels. Moreover, no individual parcels had effects stronger than what we observed at the community level.

**Figure 4.**
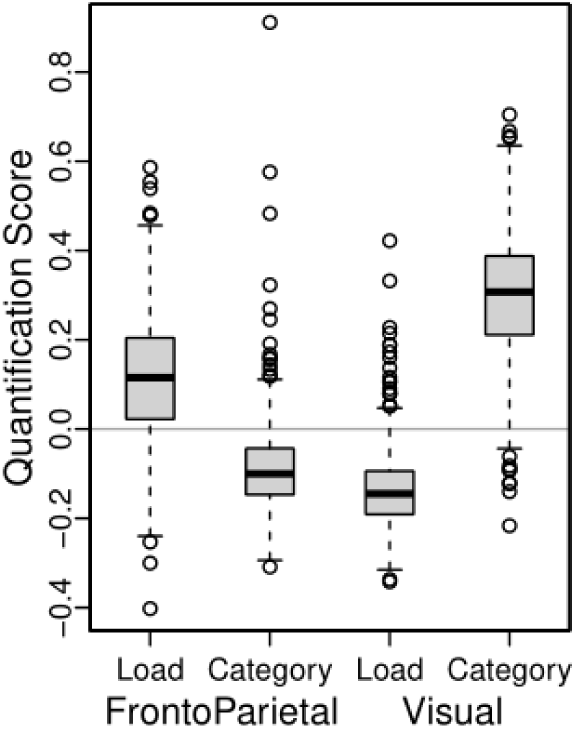
Distribution of Load and Category quantification scores for individuals’ matrices, by community. The Load quantification scores tend to be higher than Category in FrontoParietal, but the reverse in Visual. Boxplots for participants by subject group are shown in S2.3.

Although the community-based double dissociation was highly robust at the group level, there was also clear individual variation in the quantification scores. If this variability reflects functionally meaningful individual differences in brain coding of relevant task dimensions, then it should also be predictive of individual differences in behavioral task performance. To examine this question, we used N-back d’ as the behavioral measure (collapsed across Load and Category). The analysis strongly confirmed the hypothesis of functional significance, as highly selective brain-behavior relationships were observed (Figure 5). Specifically, the Load score was positively correlated in the FrontoParietal community (r=.31, p <.001), such that individuals showing a higher score (higher fidelity of load-based coding) had better N-back performance. Yet in the Visual community, the reverse pattern was present, with Load score correlating negatively with performance (r=−.19, p<.001), such that stronger Load coding predicted poorer N-back performance. On the other hand, Category scores were somewhat more weakly correlated with behavioral performance, and also showed the opposite profile (i.e., negative correlation for FrontoParietal, r=−.15, p<.001; positive correlation for Visual, r=.27, p<.001).

**Figure 5.**
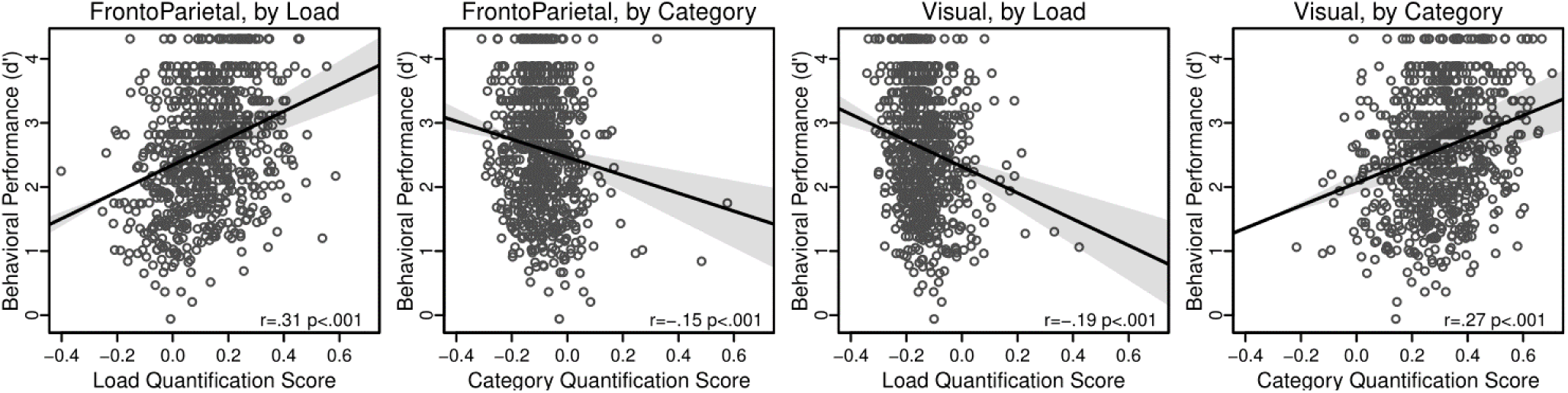
Relationship between individual behavioral performance (d’) and matrix quantification for Load and Category. Bands are .95 confidence intervals for the regression line, calculated with `lsfitci` (Wilcox, 2017). Statistics by subject group are in S2.4.

Although each of these correlations indicate high brain-behavior selectivity, they also suggest the possibility that all four scores are independently predictive of N-back performance. On the other hand, it seems likely that Load-based coding might be the most strongly associated with performance in the N-back, given the presumed dependence of the task on WM processes. To examine this issue, the data were submitted to a multiple regression analysis, with behavioral performance as the outcome variable, and all four quantification scores as potential predictor variables. This analysis confirmed that both Load scores were independently predictive of performance, with FrontoParietal Load positively predictive (beta=.32, p<.001) and Visual Load negatively predictive (beta=−.18, p=.0036). However, with all four predictors in the model, although explaining 16% of task variation, neither of the Category indices made independent contributions to predicting N-back task performance (FrontoParietal p=.4, Visual p=.1; full results in S2.5). Together, these results converge on the interpretation that individuals exhibiting strong Load coding in FrontoParietal regions will tend to have better N-back performance, while those showing strong Load coding in Visual regions will have poorer performance.

### Genetic Influences on Activation Pattern Similarity: Analyses in Pairs

After establishing the validity and utility of pattern similarity analysis for examining anatomically selective patterns of individual difference in neural coding, the second set of analyses examined the similarity of brain activation patterns in pairs of related and unrelated individuals. These analyses were conducted to provide a stronger and more novel test of the hypothesis that task-specific activation patterns (e.g., in the FPN) are genetically influenced and reflect individual differences. Adapting the approach used above, we again created pattern similarity matrices, but now computed the similarity of *paired* individuals, to quantify the degree to which activation in one member of the pair matches the other. Specifically, activation pattern similarity was again measured with Pearson correlation, but computed for all sixteen pairwise combinations of the four parameter estimate images across individuals (e.g., the 0-back Face of one twin to the 2-back Face of their co-twin; Figure 2). Example matrices for one pair of people from each group are in Figure 6; additional examples and group-level average matrices are provided in S3.1. The appearance of these pairwise matrices (Figure 6) is broadly similar to the matrices for individuals (Figure 3): highest correlations in the lower right (cells sharing 2-back Load) in FrontoParietal, but a checkerboard pattern (higher correlation for cells sharing Category) in Visual.

**Figure 6.**
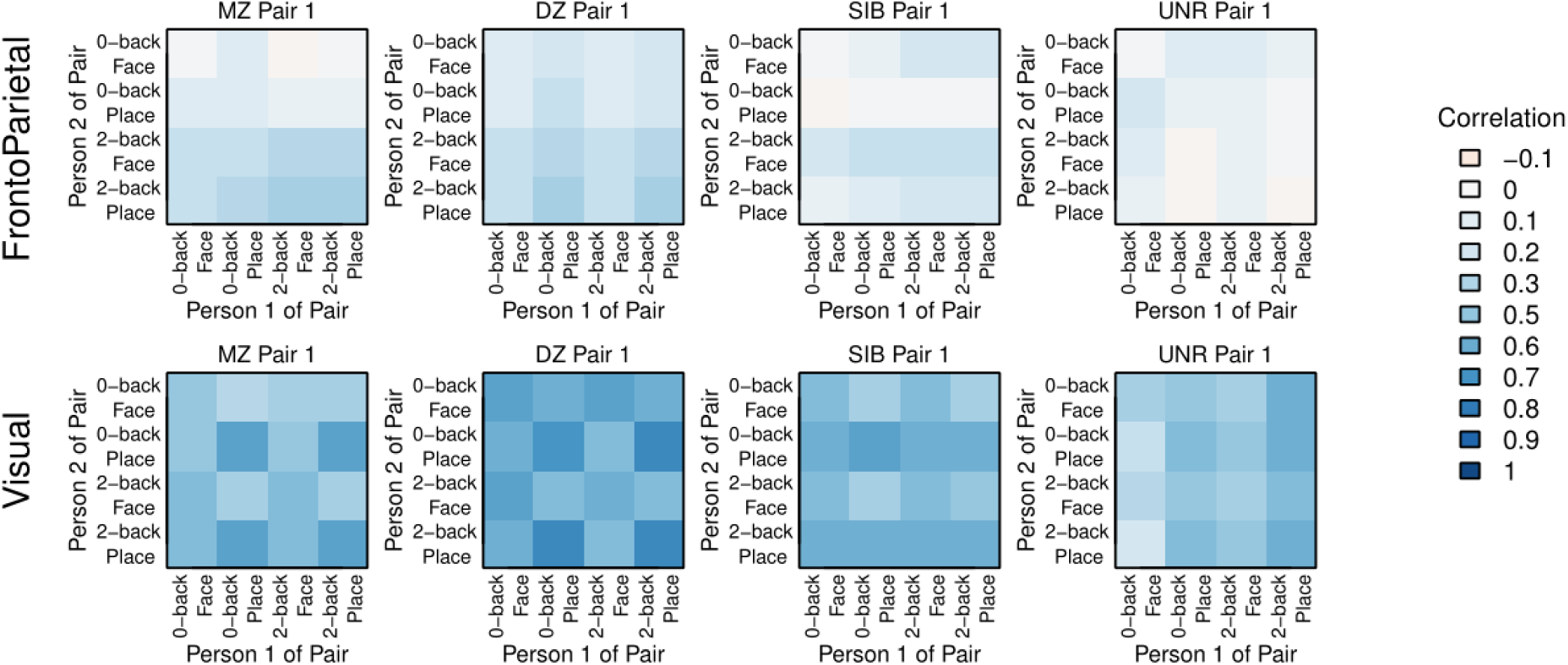
Similarity matrices for the two communities in four paired people. More examples and the group averages are in S3.1. As in the Figure 3 matrices for individuals, these pairwise similarity matrices tend to resemble the Category Reference (Figure 2) when calculated from Visual activation patterns, but the Load when calculated in FrontoParietal. The resemblance and correlations are strongest in the MZ and DZ pairs, with little similarity seen in UNR FrontoParietal.

Unlike the matrices for individuals described previously, the pairwise matrices are not symmetric, and so the diagonal is meaningful, containing the across-person within-condition correlations for each of the four matched conditions (e.g., 0-back Face in one twin with 0-back Face of their co-twin). These values along the diagonal, when contrasted across MZ, DZ, SIB and UNR, provide evidence of heritability of the condition, and parallel the type of analyses conducted by Polk et al. (2007) and Pinel et al. (2015). However, in these prior studies, analyses were restricted to visual occipitotemporal regions. Here, we were able to not only test for replication (with a much larger sample size), but also to extend these previous findings to FrontoParietal regions and to WM-related conditions. Replicating prior results, we found significantly greater Visual activation pattern similarity in MZ twins than DZ twins, non-twin siblings (SIB), and unrelated people (Figure 7; p<.001 in all pairwise t-tests, S3.2). However, unlike the prior work, we also found that the DZ and SIB pairs showed significantly higher pattern similarity than UNR pairs, even though the latter were also matched on age and gender (the significance holds even if unrelated people are matched on behavioral performance; S5.1). No significant differences between DZ and non-twin SIB pairs were found in any comparison (indicating an absence of special twin environmental effects). The stronger participant group effects in our study are likely due to the increased power and precision provided by the larger sample sizes: 105 MZ and 78 DZ twin (with an additional 99 SIB) pairs versus 11 MZ and 11 DZ twin pairs in (Polk et al., 2007) and 16 MZ and 13 DZ twin pairs in Pinel et al. (2015).

**Figure 7.**
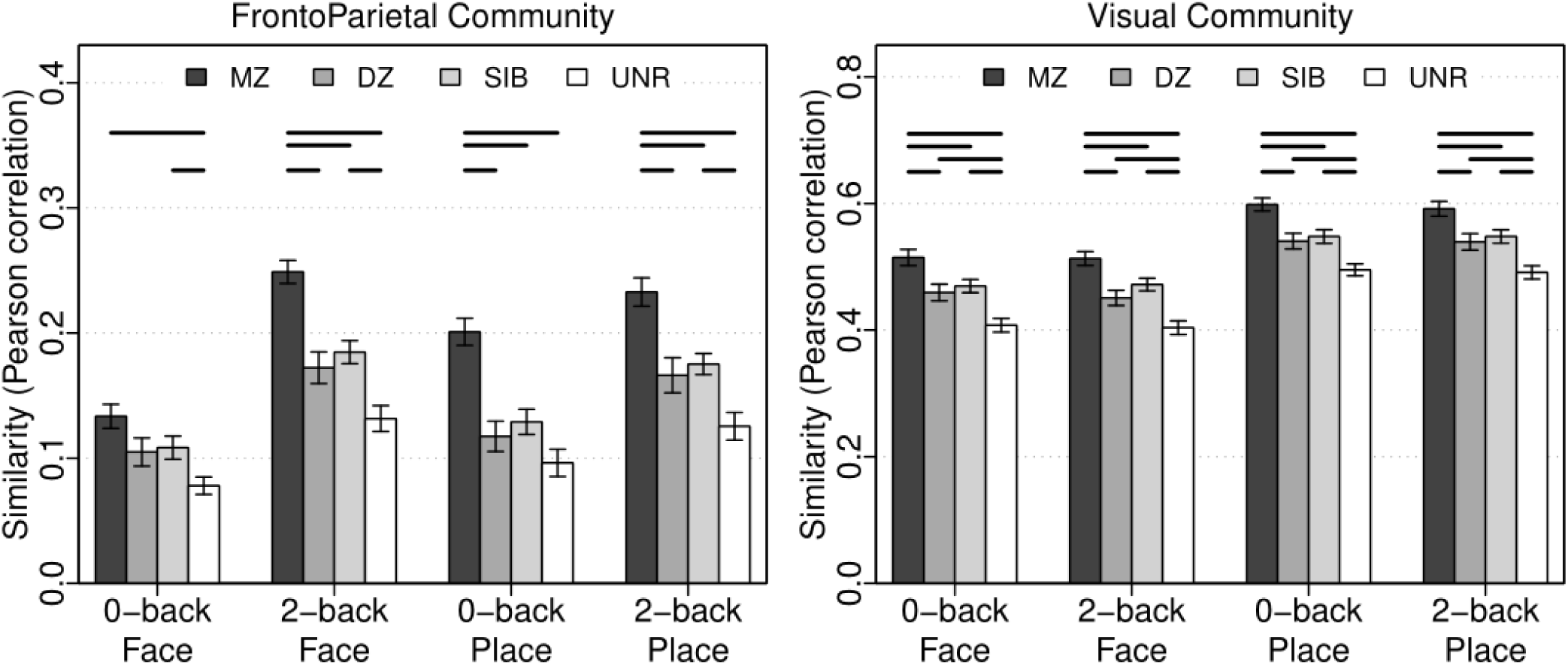
Mean similarity of matched conditions (Figure 6 matrix diagonals) in the FrontoParietal and Visual communities. Error bars are standard error of the mean (SEM). Horizontal lines indicate bars that significantly (p <.0083, Bonferroni correction of .05 for 6 comparisons) differ in a robust t-test. The full dataset and statistics are shown in S3.2.

Although Polk et al. (2007) and Pinel et al. (2015) only examined visual occipitotemporal ROIs, we carried out the analysis in the FrontoParietal community as well (Figure 7, S3.2). Parallel findings emerged: within FrontoParietal, pairwise t-tests also showed significantly greater similarity in MZ twins than DZ twins, SIB, and UNR (p<.001), and likewise greater similarity among DZ and SIB pairs relative to UNR (but again no differences between DZ and SIB). The exception to this pattern was in the 0-back Face condition, in which only the difference between MZ and UNR had p<.001. The reduced similarity in FrontoParietal for 0-back load is unsurprising, as we expected FrontoParietal activation to increase with cognitive load. In a parallel heritability analysis, shown in S3.4, we found that activation in the FrontoParietal community was more greatly attributable to individual-specific environmental effects than the Visual community, as indexed by the overall lower MZ similarity and consequent estimates of E (e^2^ ranged from .41 to .49 for Visual vs. .75 to .87 for FrontoParietal). In addition, while familial effects (genetic and common environment, estimated as 1-e^2^) were more pronounced for the Visual than FrontoParietal community, the extent to which additive genetic factors (i.e., heritability) influenced similarity in both communities was equivalent (a^2^ ranged from .05 to .16) such that the increased familial correlation in the Visual community was primarily due to stronger effects of common environment (c^2^ ranged from .38 to .41 for Visual vs. .04 to .10 for FrontoParietal).

Together, these findings replicate and extend the work of Polk et al. (2007) and Pinel et al. (2015), by demonstrating a clear role for heritable factors that are present in activation patterns not only within occipitotemporal visual regions, but also in frontoparietal regions related to working memory and executive control. Moreover, the results provide convincing evidence that FrontoParietal activation pattern similarity effects are dominated by genetic factors, with very little influence of shared environment or other confounding demographic factors (age, gender, etc.).

### Genetic, Anatomic, and Task Specificity in Pairwise Activation Similarity Patterns

Although the analyses reported above were useful for extending the findings of Polk et al. (2007) and Pinel et al. (2015), and for confirming that activation pattern similarity approaches can be used to estimate heritability effects, they do not exploit the full power of the methodology, because they only use the diagonals of the similarity matrices (correlation between pairs of individuals on matched conditions). The alternative (RSA-style) approach quantifies how well the full similarity matrix for each participant pair conforms to the reference matrices (Figure 2), and consequently, provides a more sensitive test of whether pairwise similarity is preferentially strong for a particular representational coding scheme (Load or Category). Moreover, this approach avoids the confounds inherent in correlating conditions of the same type (e.g., if two participants are found to have similar activation patterns for 2-back Place, it could be due either to the shared Load or the shared Category). Consequently, we next computed the pairwise Load and Category quantification scores in each of the two communities (Visual, FrontoParietal), for paired participants of all four types (MZ, DZ, SIB, UNR).

The pairwise Load and Category quantification scores showed a high degree of anatomic specificity. In FrontoParietal, Load scores were significantly greater than Category in all subject groups (p<.001; Figure 8 and S4.1). Conversely, in Visual, Category scores were much greater than Load in all subject groups (p<.001). Importantly, although this double dissociation is of the same form as observed in the individual analyses, it reflects an independent measure of task coding specificity. In particular, the pairwise scores reflect selective activation pattern similarity between individuals: a high Load quantification score in FrontoParietal indicates that the two individuals’ activation patterns are more similar when the WM load is the same (e.g., 2-back Face in one person and 2-back Place in their twin) than when the WM load is different (e.g., 2-back Face in one person and 0-back Face in their twin).

**Figure 8.**
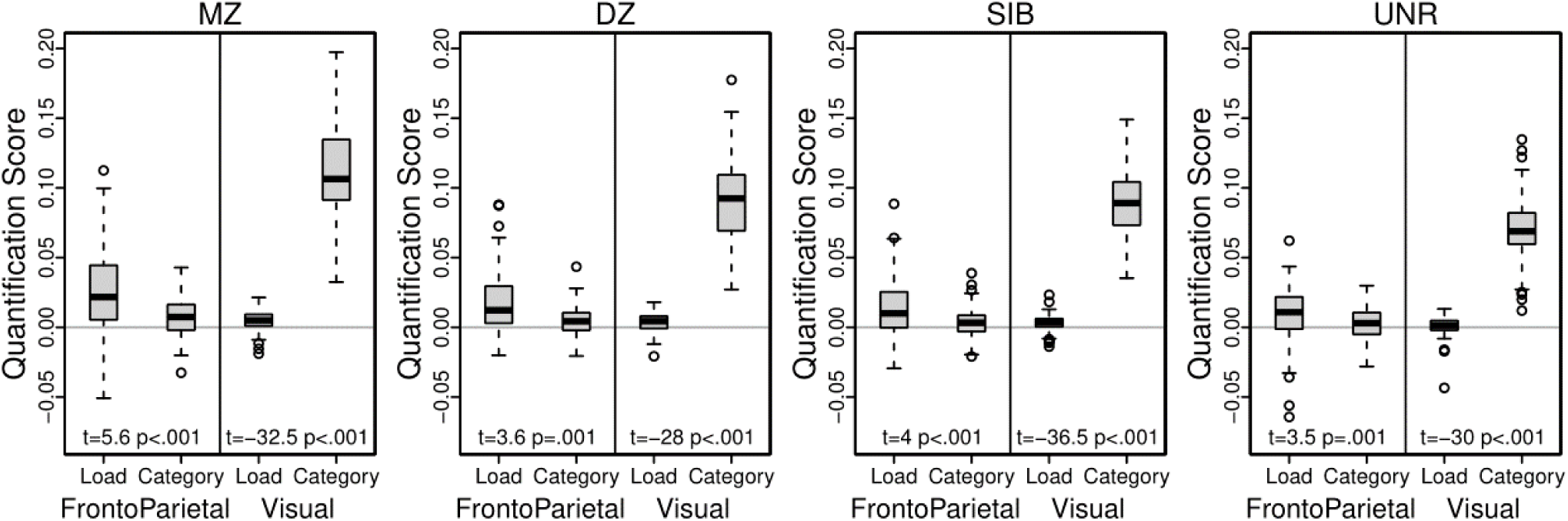
Distribution of pairwise Load and Category quantification scores, by Community. Load quantification scores are higher than Category in FrontoParietal, but lower in Visual. A robust pairwise t-test was performed within each community and subject group, as listed at the bottom of each plot (YuenTTest, trim=0.1; S4.1a). The full dataset is shown in Figure 9; one MZ FrontoParietal Category outlier at −0.18 not shown.

Moreover, this coding specificity also showed clear effects of pair group: most prominent in MZ twins, least in UNR individuals (and the same pattern of results was found even when selecting UNR pairs in which the pair members were matched on behavioral performance; S5.1b). Interestingly, these imaging results show genetic similarity effects similar to the analyses of behavioral performance. Using robust ANOVAs to test for an influence of subject group within each of the four combinations of Community and Quantification yielded highly significant effects for Load in FrontoParietal (F=5.6, p=.001; S4.1b) and Category in Visual (F=35.5, p<.001), but not for Category in FrontoParietal (F=2.2, p=.093). Using post-hoc tests to explore these significant models, Load in FrontoParietal showed not only MZ > UNR (p<.001), but also MZ > SIB (p<.001) and MZ > DZ (p=.015). Likewise, Category in Visual showed MZ greater than all three other groups (p<.001); further, DZ and SIB were significantly greater than UNR (p<.001). There was no hint of a significant difference between DZ and SIB in either model (both ps>.3), suggesting that environmental factors unique to twins have a minimal impact on the similarity of brain activation patterns. However, the DZ/SIB scores were more similar to the MZ scores than would be expected if heritable factors alone were responsible for their similarity, indicating the role of common environmental effects on the scores for both communities. Together, these results suggest that brain activation pattern similarity measures are robustly heritable, influenced by familial environmental factors, and are observed most clearly when taking into account the task coding present in the particular brain area (weaker genetic relatedness effects were found for Category in FrontoParietal and Load in Visual).

We also examined similarity effects when quantifying Load with only 2-back trials in the reference matrix (S4.1), suspecting the greater activation occurring during the high WM load condition would make activation pattern similarity in twins, if present, more pronounced (though at the possible cost of increased quantification score variance, since twelve cells go into the calculation instead of all sixteen). Indeed, the finding of Load greater than Category in FrontoParietal but Category greater than Load in Visual was also present when using only 2-back trials for Load quantification (S4.1a). The robust ANOVAs found a stronger effect of subject group in FrontoParietal with 2-back Load (F=7.1, p<.001), but no effect in Visual for 2-back Load (F=.6, p=.62, S4.1b). Conversely, if only 0-back trials are used for Load quantification there is no effect of subject group in either Community (S4.1b). An alternative interpretation of these results is that they are primarily driven by activation differences across conditions, e.g., a univariate increase in activation during the high load condition could cause an apparent increase in similarity. To investigate this possibility, we compared the mean activation in each pair of participants but found no load-related relationships (i.e., high mean FrontoParietal 2-back activation in one person does not predict high activation in their co-twin; S5.2c). Together, these results underscore the idea that these similarity measures are most clearly identifying genetic relatedness effects when the task coding dimension matches the functionality of the particular brain network.

### Brain-Behavior Relationships in Pairwise Activation Similarity Patterns

The above set of analyses confirm the presence of heritability effects in BOLD activity, while using a multivariate pattern similarity approach. A test of the functional relevance of these pairwise scores is whether they, like the individual scores, are related to behavioral performance. Figure 9 shows the quantification scores for all participant pairs, with pairs ordered by behavioral performance (mean d’). Behavioral performance clearly explains some of the variability: pairwise quantification scores tend to increase as performance increases. Note that these results again showed a high degree of specificity: in FrontoParietal, the correlation between Load and behavioral performance was significant in all related pairs (MZ, DZ, SIB, p≤.002) but was not significant for UNR pairs (p=.1); likewise, there was no association for any group when using the Category instead of the Load score (all p>.2). In contrast, within Visual, the correlation between Category and performance was significant for MZ and DZ twins (p<.005) and marginally so for SIB and UNR pairs (p<.025), but much less convincing when using Load instead of Category for quantification (Figure 9, S4.2). The anatomic and task dependent relationship with behavioral performance is even more striking when using only 2-back trials for Load quantification, but absent if only 0-back trials are used (S4.3). However, the MZ correlation was not significantly greater than the DZ or SIB correlations (which did not differ from each other: e.g. for 2-back trials for Load in the FrontoParietal community: rMZ=.46, rDZ=.49, rSIB=.43) suggesting that the relationship between behavioral performance and variability in BOLD activity might be due to common environmental rather than heritable factors.

**Figure 9.**
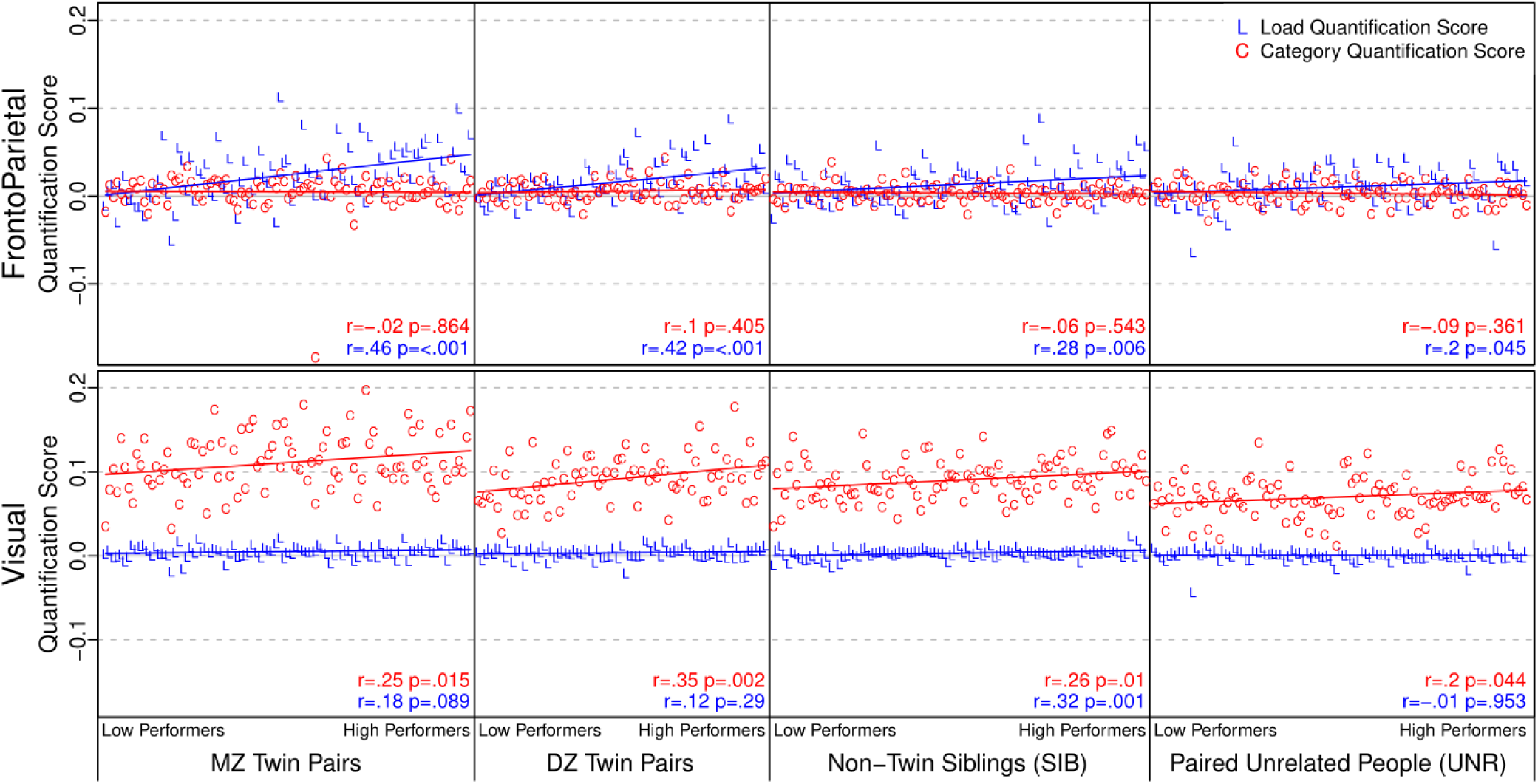
Pairwise quantification scores, arranging participant pairs along the x-axis in order of increasing behavioral performance within each subject group (d’ averaged over the two participants; pairs with missing performance for either member omitted). The best performing pair is at the right of each subject group, so higher performance is associated with higher quantification scores. Listed r and p values are from correlating quantification scores against the behavioral performance rank ordering; S4.3 and the main text give correlations against the mean d’. S4.2 has versions of this figure unsorted by behavioral performance and with different Load quantification. S5.1c has this figure for a group of unrelated participants chosen to have similar behavioral performance.

Given the similarity of these patterns to what was observed when using the individual, rather than pairwise scores, one possible concern is that the correlations between the pairwise scores and behavior are purely a reflection of shared variance with the individual scores. To test for this possibility, we conducted multiple regression analyses, predicting the pairwise score with not only behavioral performance (from each member of the pair separately) but also with the individual quantification scores from the pair members (S4.4). The results were again specific: in FrontoParietal, the pairwise Load quantification scores tended to be associated with behavioral performance, even after including the individual quantification scores as predictors. This pattern was most strongly present for MZ twins (β=.33, p<.001 for including twin 1’s d’; β=.14, p=.14 for twin 2’s d’; S4.4), but with similar trends in the DZ and SIB pairs as well. Model comparison reinforced this impression of the usefulness of including all four predictors in the multiple regression: the full model (d’ for person 1 of pair, d’ for person 2 of pair, quantification score for person 1, quantification score for person 2) outperformed the model with the individual quantification scores only (p<.001 for MZ; p=.06 for DZ; p=.098 for SIB). The difference is even more striking when only 2-back trials are used for Load quantification: the full model outperformed the quantification scores-only model at p <.001 for MZ; p=.0017 for DZ; p<.001 for SIB. The findings are dramatically different in Visual: after accounting for the individual Category quantification scores, there was no additional relationship between N-back task performance and the pairwise scores in any of the subject groups. A final control analysis (S5.2c) also included the mean (i.e., univariate) FrontoParietal and Visual activation, treating the condition difference contrasts (2-back – 0-back and Place – Face) as additional predictor variables in the multiple regression, to determine whether the above effects could be explained by the presence of mean activation differences between the conditions. Yet even with the univariate predictors included in the model, the FrontoParietal Load pairwise quantification score was still found to be associated with N-back behavioral performance, suggesting that this association could not be fully explained by pairwise similarity in load-related univariate activation levels.

Together, these findings strongly underscore the selective utility of FrontoParietal regions as functional markers of WM load-based coding and of the variability in such load-based coding both in individuals and related pairs. Thus, better N-back task performance is predicted both for individuals that show stronger evidence of selective Load coding, and additionally, for related pairs that show greater similarity in their Load coding patterns. Conversely, neither Visual regions nor variation in stimulus-based coding can serve as equivalent predictors of performance, which again reinforces the selective importance of FrontoParietal Load coding to N-back performance.

## Discussion

The primary goal of this study was to test whether multivariate pattern similarity approaches could provide increased sensitivity and leverage for revealing the neural coding properties, individual differences, and genetic similarity effects present within the frontoparietal network (FPN). In this regard, the results provide compelling support along four different dimensions. First, we found clear evidence of *functional anatomic specialization*, such that while visual occipitotemporal cortex was selectively sensitive to similarity effects related to perceptual category (face, place), the FPN was selectively sensitive to similarity effects related to working memory (WM) load, a higher-order cognitive dimension strongly related to executive function and cognitive control. Second, pattern similarity indices in FPN showed clear evidence of *systematic individual variation*, and moreover, these individual differences were strongly associated with task performance, such that individuals exhibiting stronger selectivity to WM load coding also performed better on the N-back task. Third, we found that pattern similarity indices could be used to clearly reveal a gradient of *genetic relatedness*, such that identical (MZ) twins showed the strongest levels of selective pattern similarity to WM load in the FPN, with lower, but still significant degrees of similarity found among pairs showing 50% genetic relatedness (i.e., fraternal/DZ twins and siblings). Finally, we identified a new metric for *quantifying “pairwise” variation*, in that genetically related pairs showing greater degrees of selective similarity for WM load coding in FPN also had uniquely better N-back performance. Taken together, the results strongly reinforce the coding specificity principle, in demonstrating that FPN shows unique coding properties that are both sensitive to multiple dimensions of variation (individual, genetic) and also functionally relevant for task performance. We next describe further implications of the present results, as well as their relationship to prior work.

### Neural Coding of WM Load in FPN

The current results are consistent with a large neuroimaging literature indicating the importance of the FPN in WM and executive control functions (Braver & Ruge, 2006; Niendam et al., 2012; Rottschy et al., 2012). However, the current work extends beyond much of this prior literature, which has tended to rely on univariate measures of FPN involvement in WM. Indeed, in prior work, the focus has typically been on demonstrating increased or decreased FPN activity as a function of WM load, or other relevant variables, such as the type of information being maintained, updating or manipulation requirements, and distractor-related interference. By contrast, multivariate approaches shift the focus to the *pattern of activity*, potentially providing greater traction regarding how WM load is *represented* in the FPN. In particular, multivariate approaches can provide information regarding the WM-related content being coded by a region, even when the mean (i.e., univariate) level of activity may not change or be sufficiently sensitive (Harrison & Tong, 2009; Riggall & Postle, 2012; Serences, Ester, Vogel, & Awh, 2009). In the current study, we specifically employed multivariate pattern similarity techniques to demonstrate that the structure of activation similarity or dissimilarity across WM conditions can also be informative. For example, here we demonstrated that FPN regions show significantly greater activation similarity in conditions that share the same WM load, even when the content of information being maintained can change, relative to posterior occipitotemporal regions. Moreover, we established that such similarity metrics are functionally important, in that they may reflect how well WM load information is represented in an individual (or pair of individuals), as this information appears to predict more accurate task performance.

Although the current study highlights the potential of pattern similarity approaches for testing questions regarding WM, executive control, and FPN function, it should be clear that this work represents just an initial step, and indeed the questions being asked in the current study were cast at a relatively coarse grain. For example, the current study focused on just the N-back task, with only two levels of WM load, and relied on block-related measures of activity. However, because pattern similarity analyses are eminently flexible, the approach could be easily extended to compare various WM task paradigms, to focus on different load levels, or to utilize event-related designs, which would enable a more fine-grained focus on various within-trial events (encoding, delay, probe decisions) and/or activity dynamics within the trial (King & Dehaene, 2014). Such extensions are likely to be highly fruitful, and could be used to resolve important questions raised by the current work, such as the finding that the similarity structure of the 0-back seemed different from the 2-back. By examining other load levels (e.g., 1-back, 3-back) it could be better determined whether there are qualitative differences that make some load levels more distinct from others (e.g., 2-back and 3-back may be more similar to each other than they are to 0-back or 1-back). Indeed, although there is a growing literature utilizing MVPA decoding approaches within the domain of WM and cognitive control (D’Esposito & Postle, 2015), the use of pattern similarity measures is still sparse in this domain, relative to its adoption in other cognitive domains, such as perceptual coding (Chikazoe, Lee, Kriegeskorte, & Anderson, 2014; Kriegeskorte et al., 2008) and episodic memory (Dimsdale-Zucker & Ranganath, 2018; LaRocque et al., 2013; Xue et al., 2010). We hope that the utility of the pattern similarity approach demonstrated here will encourage other researchers to begin applying it to a broader range of questions in WM and executive control.

### Neuroimaging of Individual Differences

Within cognitive neuroscience there has been steadily increasing interest in using neural measures to better capture and characterize individual differences (Braver, Cole, & Yarkoni, 2010; Cooper, Jackson, Barch, & Braver, 2019; Gordon et al., 2017; Gratton et al., 2018; Satterthwaite, Xia, & Bassett, 2018). This focus on individual differences has partly been driven by the advent of large-scale neuroimaging studies, which are optimized for sensitivity to detect reliable individual variation (Cooper et al., 2019). Indeed, a primary rationale and goal of the HCP was to define individual variation in the human connectome (Van Essen et al., 2012). The success the HCP and others like it have led to a great deal of excitement around the concepts of personalized neuroscience (Satterthwaite et al., 2018) and “connectome fingerprinting” (Finn et al., 2015). Yet much of the recent excitement around individual difference-focused datasets such as the HCP, has been on characterizing individual differences from resting-state functional connectivity (rsFC), and under task-free states (Gratton et al., 2018; Tavor et al., 2016) rather than on task-based fMRI activation patterns.

The current findings illustrate some of the unique advantages of task-based fMRI patterns in terms of detecting functional and anatomic specificity of individual variation. In particular, a key finding was that individual differences were found to be dependent on task context. Within the FPN, the individual differences in activation patterns that predicted task performance were selective to coding of WM load; individual differences in the coding of perceptual category in FPN had no relationship to task performance. Conversely, when looking at visual regions, the strength of WM load coding negatively predicted performance, such that individuals with strong coding of WM load in Visual tended to have poorer task performance. These context-specific individual differences patterns also highlight the utility of pattern similarity approaches for understanding the nature of individual variation. The findings reinforce the notion that it is the coding specificity of FPN and visual regions that is functionally critical for optimal task performance. In other words, the findings demonstrate that it was the individuals showing the strongest functional specificity – coding WM load only in FPN and perceptual category only in visual regions – who exhibited the best performance. This relationship should only be present if coding specificity is functionally relevant for task performance. Together, this work suggests that computing pattern similarity-based quantification scores that compare alternate coding schemes could be a powerful approach for revealing individual differences. Future work is needed, though, to demonstrate that such approaches could also work well in other domains. For example, in the HCP Gambling task, quantification scores could be computed to identify reward or punishment coding and determine whether individual differences in coding scores might predict functionally relevant behavioral indices (e.g., trait reward or punishment sensitivity).

### Genetic Relatedness Effects

The current approach represents a departure from the standard methods used in genetic neuroimaging analyses, in which univariate measures of ROI or voxel-based activation contrasts are tested for genetic correlation and subsequent statistical modeling of heritability. Instead, the method used here to identify potential genetic relatedness was one that harnessed potentially more powerful multivariate pattern similarity approaches. Although such approaches have rarely been used in this literature, they may be particularly well-suited for analyses of genetic relatedness. The key hypothesis is that, if influenced by genetic factors, then activation pattern similarity in paired individuals should track their degree of genetic similarity (i.e., relatedness: MZ > DZ and non-twin siblings > unrelated). The utility of the pattern similarity approach for identifying heritability in brain activation was first demonstrated by Polk et al. (2007), in a study focused on perceptual category coding in visual regions, with similar findings obtained by Pinel et al., (2015).

The current findings replicate this earlier work, but also extend it in important ways. Specifically, by harnessing the large sample size of the HCP, we were able to confirm the robustness of heritability effects, with clear evidence of MZ > DZ in both Visual and FrontoParietal. However, due to our considerably larger sample size and ability to combine data on DZ twins and non-twin SIBs, we were able to obtain a more precise estimate of common environmental influence. While our MZ correlations were similar to those reported by Pinel et al., (2015), our DZ and SIB correlations were nearly double their estimate, resulting in fairly robust estimates of common environment (see extended discussion in S3.5). Thus, even though our estimates of individual-specific environment approximate those reported by Pinel et al. (2015), familial similarity for Visual was due to genetic and common environment in our study. In contrast, Polk et al. (2007) reported higher MZ and DZ similarity than the current study or the work of Pinel et al. (2015), supporting the role of common environmental effects in addition to heritable influences, but underestimating the role of individual-specific environmental factors.

Potentially the most important methodological advance of our work over by Polk et al. (2007) and Pinel et al. (2015) is that we evaluated pattern similarity effects in paired individuals across a full set of task conditions, rather than restricting analyses to matched conditions. This extension of the pattern similarity approach enables construction of a full *similarity matrix*, similar to the representational similarity analysis (RSA) popularized by Kriegeskorte et al. (2008). In this approach the observed similarity matrix can be compared against theoretically-specified reference matrices to compute quantification scores, which can then be used to test between alternative coding models. Importantly, quantification scores incorporate the similarity of the twin pair members when they are performing the same task (as is usual, such as the estimation of cross-pair within-task correlations in genetic modeling (Neale & Maes, 2002)), but also the similarity when they are performing different tasks (e.g., one pair member performing 2-back Face and the other performing 2-back Place; cross-pair cross-task correlations).

The RSA-style quantification score approach may be a more reliable and sensitive way of revealing familial effects than even found in the prior studies adopting pattern similarity analyses. The power of this approach was most clearly demonstrated in the direct comparison of genetic relatedness influences on pairwise quantification scores in Visual and FrontoParietal, as these clearly indicated the specificity of observed heritability effects (Figure 8). In particular, although Visual quantification scores showed evidence for heritability in perceptual coding (i.e., scores showing a MZ > DZ, SIB > UNR pattern), these regions showed no evidence of heritability with regard to WM coding. Conversely, FrontoParietal quantification scores indicated heritability effects on WM coding, replicating prior results (Blokland et al., 2008, 2011), but showed no evidence for heritability with regard to perceptual coding. Thus, the current findings make a stronger case for the functional-anatomic specificity of heritability effects than has been observed in prior genetic neuroimaging studies examining the FPN. Such findings of functional-anatomic specificity would not be possible in genetically-informed studies that are solely focused on task-free states or do not manipulate task context.

### Quantifying “Pairwise” Variation

A novel advantage of the pattern similarity approach used here is that it produces a unique quantification score for each twin pair, rather than treating each twin as an individual observation. We demonstrated that this pairwise measure is functionally important, in that it was reliably associated with task performance. Critically, that the pair-related predictive effects were unique; i.e., over and above the variance explained by individual quantification scores. While not implemented in the current analysis, it would even be possible to estimate the statistical significance of the quantification score for a given twin pair by creating null distributions from the estimated similarity of each member with unrelated members of the dataset (e.g., comparing the differences in pairwise Load quantification scores when a given individual is paired with their co-twin relative to when they are paired to a set of unrelated individuals).

The brain-behavior relationships uncovered using the pairwise quantification scores are compelling and provocative in their implications. Specifically, we observed that genetically related pairs showing more similarity to each other (greater FrontoParietal Load quantification), also tended to show better N-back task performance. However, this observation leads to an additional question: why might such a pattern be present? Although our interpretation remains speculative, we suggest that it might be due to the combination of two factors: 1) “sharper” or higher fidelity task coding patterns in high performing individuals; and 2) increased anatomic or functional similarity among related individuals. With regard to the first factor, it is generally assumed that people with higher task performance are more focused and engaged with the task, and our own prior work provides initial evidence that stronger and more distinct task coding patterns would be expected in these individuals (Etzel et al., 2016). If we assume this first factor to be correct, we might then also expect that high performing individuals would have activation patterns that would tend to be similar (at the vertex level) to their twin or non-twin sibling. In particular, we speculate that genetically related individuals would be more likely to show similar activation patterns to each other when both are coding task-relevant variables, such as WM load, in an optimal (i.e., veridical) manner. Logically, there are far more ways to perform a task poorly (e.g., not attending, forgetting the stimulus, confusion about the instructions) than there are to perform it well. Thus, it is more likely that a pair of related individuals will show high similarity to each other when both have an optimal coding of WM load. Note that we are not claiming that related individuals (or twins) are more likely to show higher fidelity or less noisy task coding patterns in general, but that similarity of activation patterns can be used as an additional way to identify individuals that are likely to show stronger coding of relevant task variables.

It is also noteworthy that the relationship between behavioral performance and activation similarity in MZ twin pairs did not significantly differ from DZ pairs or SIBs, suggesting that while familial effects play a role, they are more likely to be of an environmental nature. Under the equal environments assumption (Robert Plomin, Willerman, & Loehlin, 1976), MZ and DZ twins (and in our case, SIBs) share some environmental factors to the same extent. Our initial twin analyses of d’ did not support the role of shared environment (S1.6), but it is plausible that the detection of shared environment, while underpowered in the univariate model, was better estimated when examined in the context of brain-behavior relationships. Such common environmental influences may reflect the impact of socioeconomic status and parental educational achievement, which are known to impact executive functioning (Hackman & Farah, 2009; Noble et al., 2015), and which may have resulted in twins (both MZ and DZ) and siblings being exposed to similar educational opportunities. As academic achievement is heritable (Cesarini & Visscher, 2017), common environmental estimates in such instances may be upwardly biased in the presence of undetected positive and passive gene-common environment covariance (rAC; i.e., parental educational achievement and executive functioning creates the educational environment that the twins are passively exposed to via neighborhood and school choice (Verhulst & Hatemi, 2013)). Undetected assortative mating can also inflate estimates of common environment and there is considerable support for primary assortment for intelligence (Coventry & Keller, 2005; R. Plomin & Deary, 2015). Alternatively, we might posit that, given the complexity of these multivariate correlational indices, we were underpowered to tease apart genetic and environmental sources of familial similarity. Another possibility is that the equal environments assumption is not met, but a sensible test of that hypothesis was not possible in this sample of adult twins (which was still of small size for such genetic modeling), particularly given the potentially complex multivariate relationships (e.g., parsing these relationships in twins whose self-reported zygosity diverged from their genomically-determined zygosity).

### Limitations and Future Directions

In the current work we purposely restricted the primary analyses to two large communities (FrontoParietal and Visual) and a 2×2 condition subset (0-back/2-back × Face/Place) of a single task (the N-back), to allow for clear predictions regarding the functional specialization of each brain region. We were thus able to establish the validity and utility of activation pattern similarity approaches, which were then more fully investigated using novel pairwise analyses. However, the clarity gained with these restrictions necessarily means that the findings were limited to the two communities investigated, rather than sampling the whole brain. Likewise, by restricting the analysis to whole communities, rather than the individual parcels of which they are composed, the results provide information related only to a very macro level of brain organization. In supplemental analyses, we took an initial step towards addressing these limitations, by conducting a whole-brain investigation of effects at the level of individual parcels. These analyses were largely supportive and convergent with the primary results. In particular, they confirmed that perceptual category coding was almost exclusively restricted to occipitotemporal cortex parcels within the Visual community. In contrast, working memory load coding was more widely distributed, but confirmed to be present in individual parcels of the FrontoParietal community, with additional strong coding found in parcels assigned to the Dorsal Attention and Default Mode communities. However, the parcel-level results also supported the choice of focusing our primary analysis at the macro level, as no individual parcel showed effects that were stronger than the two communities. Although it was outside of the scope of the current work to explore the optimal spatial scale and distributed vs. focal nature of observed activation pattern similarity effects, a more systematic investigation of this issue would seem to be a valuable direction for future work.

A second purposeful restriction of focus in the current study was to investigate the validity and utility of multivariate pattern similarity approaches for addressing questions of functional specialization, individual differences, and genetic similarity in the FPN, but without directly benchmarking these approaches against more standard univariate (i.e., activation level) analyses. Thus, it is important to be clear that we do not make strong claims regarding the superiority of multivariate approaches relative to univariate ones, only that they appear to have important potential that heretofore has not been fully exploited. Indeed, many questions remain as to the exact relationship of multivariate and univariate findings. Although fully exploring these relationships was beyond the scope of the current study, we did conduct some initial control analyses, to address a potential alternative interpretation of the results, which was that the observed genetic pair-based similarity effects and brain-behavior relationships could be fully explained by univariate activation differences among relevant task conditions in the FPN (e.g., 2-back > 0-back activity). Our analyses do convincingly rule out this explanation, since at the spatial scale of FrontoParietal and Visual communities, univariate activation levels were not significantly correlated among genetically related pairs. Likewise, even when including univariate (i.e., mean-ROI) activation differences among task conditions in the regression model, pattern similarity in related pairs was still uniquely associated with N-back task performance. Nevertheless, additional work needs to be done to better understand the boundary conditions under which univariate activity levels might contribute to multivariate pattern similarity effects, or when each technique may be preferable.

As mentioned above, even within the HCP there are a larger set of tasks beyond the N-back from which to explore individual differences and genetic effects. Based on the coding specificity principle, we would expect that these effects would each show a distinctive pattern of anatomic localization that is task context dependent. In other words, the coding specificity principle would suggest that FPN regions would only be sensitive to individual differences and genetic effects in task contexts related to WM and executive control (e.g., Relational Processing), but not in far different contexts (e.g., Motor, Emotion).

Additionally, with the exception of the behavioral data, for which we used ACE models, we computed quantitative estimates of heritability using simple adaptations of the Falconer equations. It seems possible that more sophisticated statistical analyses could be used to estimate heritability components using the full variance-covariance structure of the activation similarity matrices, along with other techniques, such as structural equation modeling. In particular, the estimates of cross-task cross-pair correlations (e.g., 2-back Face and 2-back Place compared to 2-back Face and 0-back Place) could provide a stronger basis for disentangling genetic from environmental contributions to activation patterns. Thus, another target for future work would be to more fully integrate pattern similarity analyses into heritability models.

The use of pairwise quantification scores opens the door for other types of genetic analyses. For example, one could examine the role of individual genome-wide significant variants and polygenic liability scores to WM function in quantification scores. Another possibility would be to contrast concordant and discordant MZ twin pairs (i.e., high vs. low pairwise quantification scores) to identify the role of individual-specific environmental factors, such as epigenetic signatures. Indeed, this work suggests a potential subdomain of genetic neuroimaging focused at the level of individual twin pairs, using quantification scores as a functionally relevant biomarker metric with which to subdivide and classify pairs. However, much still needs to be learned regarding the sensitivity of the quantification scores to the many different factors that can impact the BOLD signal (e.g., physiological, task, acquisition), for which simulation and modeling may prove valuable. We suspect that the quantification score approach may be especially rich for disentangling these different contributors, such as by describing characteristic pairwise similarity matrices (e.g., universally high correlation may indicate a dominant non-neural contribution). Yet even given this acknowledged early state of knowledge regarding the sources of activation pattern similarity, the results presented here highlight the utility and promise of adopting such approaches for the investigation of task coding properties, individual differences, and genetic effects within the FPN and other brain regions. We hope that future research will extend the current investigations in fruitful directions.

## Acknowledgments

This research was supported by NIH R37MH066078 to Todd Braver. Ya’el Courtney was supported by the BP-ENDURE Neuroscience Pipeline Program, NINDS grant R25NS090985. Data were provided by the Human Connectome Project, WU-Minn Consortium (Principal Investigators: David Van Essen and Kamil Ugurbil; 1U54MH091657) funded by the 16 NIH Institutes and Centers that support the NIH Blueprint for Neuroscience Research; and by the McDonnell Center for Systems Neuroscience at Washington University. Arpana Agrawal acknowledges support from NIDA grant K02DA032573.

## References

Assem, M., Glasser, M. F., Van Essen, D. C., & Duncan, J. (2019). A Domain-general Cognitive Core defined in Multimodally Parcellated Human Cortex. BioRxiv, 517599. https://doi.org/10.1101/517599

Barch, D. M., Burgess, G. C., Harms, M. P., Petersen, S. E., Schlaggar, B. L., Corbetta, M., … Van Essen, D. C. (2013). Function in the human connectome: Task-fMRI and individual differences in behavior. Neuroimage, 80, 169–189. https://doi.org/http://dx.doi.org/10.1016/j.neuroimage.2013.05.033

Blokland, G. A. M., McMahon, K. L., Hoffman, J., Zhu, G., Meredith, M., Martin, N. G., … Wright, M. J. (2008). Quantifying the heritability of task-related brain activation and performance during the N-back working memory task: A twin fMRI study. Biological Psychology. https://doi.org/10.1016/j.biopsycho.2008.03.006

Blokland, G. A. M., McMahon, K. L., Thompson, P. M., Martin, N. G., de Zubicaray, G. I., & Wright, M. J. (2011). Heritability of Working Memory Brain Activation. Journal of Neuroscience, 31(30), 10882–10890. https://doi.org/10.1523/JNEUROSCI.5334-10.2011

Blokland, G. A. M., Wallace, A. K., Hansell, N. K., Thompson, P. M., Hickie, I. B., Montgomery, G. W., … Wright, M. J. (2017). Genome-wide association study of working memory brain activation. International Journal of Psychophysiology, 115, 98–111. https://doi.org/10.1016/j.ijpsycho.2016.09.010

Bode, S., & Haynes, J.-D. (2009). Decoding sequential stages of task preparation in the human brain. Neuroimage, 45(2), 606–613.

Braver, T. S., Cole, M. W., & Yarkoni, T. (2010). Vive les differences! Individual variation in neural mechanisms of executive control. Current Opinion in Neurobiology, 20(2), 242–250. https://doi.org/10.1016/j.conb.2010.03.002

Braver, T. S., & Ruge, H. (2006). No Title. In R. Cabeza & A. Kingstone (Eds.), Cognitive neuroscience. Handbook of functional neuroimaging of cognition (2nd ed., pp. 307–348). MIT Press, Cambridge, MA.

Cesarini, D., & Visscher, P. M. (2017). Genetics and educational attainment. Npj Science of Learning, 2(1), 4. https://doi.org/10.1038/s41539-017-0005-6

Chikazoe, J., Lee, D. H., Kriegeskorte, N., & Anderson, A. K. (2014). Population coding of affect across stimuli, modalities and individuals. Nature Neuroscience, 17, 1114–1122.

Cole, M. W., Etzel, J. A., Zacks, J. M., Schneider, W., & Braver, T. S. (2011). Rapid transfer of abstract rules to novel contexts in human lateral prefrontal cortex. Frontiers in Human Neuroscience, 5, 1–13. https://doi.org/10.3389/fnhum.2011.00142

Cole, M. W., Reynolds, J. R., Power, J. D., Repovs, G., Anticevic, A., & Braver, T. S. (2013). Multi-task connectivity reveals flexible hubs for adaptive task control. Nature Neuroscience, 16, 1348–1355. https://doi.org/https://doi.org/10.1038/nn.3470

Cole, M. W., & Schneider, W. (2007). The cognitive control network: Integrated cortical regions with dissociable functions. Neuroimage, 37(1), 343–360. https://doi.org/http://dx.doi.org/10.1016/j.neuroimage.2007.03.071

Cole, M. W., Yarkoni, T., Repovs, G., Anticevic, A., & Braver, T. S. (2012). Global Connectivity of Prefrontal Cortex Predicts Cognitive Control and Intelligence. Journal of Neuroscience, 32(26), 8988–8999. https://doi.org/10.1523/jneurosci.0536-12.2012

Cooper, S. R., Jackson, J. J., Barch, D. M., & Braver, T. S. (2019). Neuroimaging of individual differences: A latent variable modeling perspective. Neuroscience and Biobehavioral Reviews, 98, 29–46. https://doi.org/10.1016/j.neubiorev.2018.12.022

Coventry, W. L., & Keller, M. C. (2005). Estimating the Extent of Parameter Bias in the Classical Twin Design: A Comparison of Parameter Estimates From Extended Twin-Family and Classical Twin Designs. Twin Research and Human Genetics, 8(3), 214–223. https://doi.org/DOI:10.1375/twin.8.3.214

D’Esposito, M., & Postle, B. R. (2015). The Cognitive Neuroscience of Working Memory. Annual Review of Psychology, 66(1), 115–142. https://doi.org/10.1146/annurev-psych-010814-015031

Dimsdale-Zucker, H. R., & Ranganath, C. (2018). Chapter 27 – Representational Similarity Analyses: A Practical Guide for Functional MRI Applications. In D. B. T.-H. of B. N. Manahan-Vaughan (Ed.), Handbook of In Vivo Neural Plasticity Techniques (Vol. 28, pp. 509–525). Elsevier. https://doi.org/https://doi.org/10.1016/B978-0-12-812028-6.00027-6

Duncan, J. (2001). An adaptive coding model of neural function in prefrontal cortex. Nature Reviews Neuroscience, 2, 820–829. https://doi.org/10.1038/35097575

Duncan, J. (2010). The multiple-demand (MD) system of the primate brain: mental programs for intelligent behaviour. Trends In Cognitive Sciences, 14(4), 172–179. https://doi.org/http://dx.doi.org/10.1016/j.tics.2010.01.004

Etzel, J. A., Cole, M. W., Zacks, J. M., Kay, K. N., & Braver, T. S. (2016). Reward Motivation Enhances Task Coding in Frontoparietal Cortex. Cerebral Cortex, 26(4). https://doi.org/10.1093/cercor/bhu327

Evans, D. M., Gillespie, N. A., & Martin, N. G. (2002). Biometrical genetics. Biological Psychology, 61(1), 33–51. https://doi.org/https://doi.org/10.1016/S0301-0511(02)00051-0

Finn, E. S., Shen, X., Scheinost, D., Rosenberg, M. D., Huang, J., Chun, M. M., … Constable, R. T. (2015). Functional connectome fingerprinting: Identifying individuals using patterns of brain connectivity. Nature Neuroscience, 18, 1664–1671. https://doi.org/10.1038/nn.4135

Glasser, M. F., Sotiropoulos, S. N., Wilson, J. A., Coalson, T. S., Fischl, B., Andersson, J. L., … Jenkinson, M. (2013). The minimal preprocessing pipelines for the Human Connectome Project. Neuroimage, 80, 105–124. https://doi.org/http://dx.doi.org/10.1016/j.neuroimage.2013.04.127

Gordon, E. M., Laumann, T. O., Adeyemo, B., Huckins, J. F., Kelley, W. M., & Petersen, S. E. (2016). Generation and Evaluation of a Cortical Area Parcellation from Resting-State Correlations. Cerebral Cortex, 26(1), 288–303. https://doi.org/doi:10.1093/cercor/bhu239

Gordon, E. M., Laumann, T. O., Gilmore, A. W., Newbold, D. J., Greene, D. J., Berg, J. J., … Dosenbach, N. U. F. (2017). Precision Functional Mapping of Individual Human Brains. Neuron, 95(4), 791–807.e7. https://doi.org/https://doi.org/10.1016/j.neuron.2017.07.011

Gratton, C., Laumann, T. O., Nielsen, A. N., Greene, D. J., Gordon, E. M., Gilmore, A. W., … Petersen, S. E. (2018). Functional Brain Networks Are Dominated by Stable Group and Individual Factors, Not Cognitive or Daily Variation. Neuron, 98(2), 243–245. https://doi.org/10.1016/j.neuron.2018.03.035

Hackman, D. A., & Farah, M. J. (2009). Socioeconomic status and the developing brain. Trends in Cognitive Sciences, 13(2), 65–73. https://doi.org/https://doi.org/10.1016/j.tics.2008.11.003

Harrison, S. A., & Tong, F. (2009). Decoding reveals the contents of visual working memory in early visual areas. Nature, 458, 632–635. https://doi.org/10.1038/nature07832

Hautus, M. J. (1995). Corrections for extreme proportions and their biasing effects on estimated values of d′. Behavior Research Methods, Instruments, & Computers, 27(1), 46–51. https://doi.org/10.3758/BF03203619

Haxby, J. V, Gobbini, M. I., Furey, M. L., Ishai, A., Schouten, J. L., & Pietrini, P. (2001). Distributed and Overlapping Representations of Faces and Objects in Ventral Temporal Cortex. Science, 293(5539), 2425–2430. https://doi.org/10.1126/science.1063736

Hendriks, M. H. A., Daniels, N., Pegado, F., & de Beeck, H. P. O. (2017). The effect of spatial smoothing on representational similarity in a simple motor paradigm. Frontiers in Neurology. https://doi.org/10.3389/fneur.2017.00222

Jung, R. E., & Haier, R. J. (2007). The Parieto-Frontal Integration Theory (P-FIT) of intelligence: Converging neuroimaging evidence. Behavioral and Brain Sciences, 30(2), 135–154. https://doi.org/DOI:10.1017/S0140525X07001185

Kane, M. J., & Engle, R. W. (2002). The role of prefrontal cortex in working-memory capacity, executive attention, and general fluid intelligence: An individual-differences perspective. Psychonomic Bulletin & Review, 9(4), 637–671. https://doi.org/10.3758/bf03196323

King, J. R., & Dehaene, S. (2014). Characterizing the dynamics of mental representations: The temporal generalization method. Trends in Cognitive Sciences, 18(4), 203–210. https://doi.org/10.1016/j.tics.2014.01.002

Koten, J. W., Wood, G., Hagoort, P., Goebel, R., Propping, P., Willmes, K., & Boomsma, D. I. (2009). Genetic Contribution to Variation in Cognitive Function: An fMRI Study in Twins. Science, 323(5922), 1737–1740. https://doi.org/10.1126/science.1167371

Kriegeskorte, N., Goebel, R., & Bandettini, P. (2006). Information-based functional brain mapping. PNAS, 103(10), 3863–3868.

Kriegeskorte, N., Mur, M., & Bandettini, P. A. (2008). Representational similarity analysis—connecting the branches of systems neuroscience. Frontiers in Systems Neuroscience, 2, 1–28. https://doi.org/10.3389/neuro.06.004.2008

LaRocque, K. F., Smith, M. E., Carr, V. A., Witthoft, N., Grill-Spector, K., & Wagner, A. D. (2013). Global Similarity and Pattern Separation in the Human Medial Temporal Lobe Predict Subsequent Memory. The Journal of Neuroscience, 33(13), 5466–5474. https://doi.org/10.1523/JNEUROSCI.4293-12.2013

Lee, J., & Geng, J. J. (2017). Idiosyncratic Patterns of Representational Similarity in Prefrontal Cortex Predict Attentional Performance. The Journal of Neuroscience, 37(5), 1257–1268. https://doi.org/10.1523/JNEUROSCI.1407-16.2016

Matthews, S. C., Simmons, A. N., Strigo, I., Jang, K., Stein, M. B., & Paulus, M. P. (2007). Heritability of anterior cingulate response to conflict: An fMRI study in female twins. NeuroImage, 38(1), 223–227. https://doi.org/https://doi.org/10.1016/j.neuroimage.2007.07.015

Neale, M. C., & Maes, H. H. M. (2002). Methodology for Genetic Studies of Twins and Families. Dordrecht, The Netherlands: Kluwer Academic Publishers B.V.

Niendam, T. A., Laird, A. R., Ray, K. L., Dean, Y. M., Glahn, D. C., & Carter, C. S. (2012). Meta-analytic evidence for a superordinate cognitive control network subserving diverse executive functions. Cognitive, Affective, & Behavioral Neuroscience, 12(2), 241–268. https://doi.org/10.3758/s13415-011-0083-5

Nili, H., Wingfield, C., Walther, A., Su, L., Marslen-Wilson, W., & Kriegeskorte, N. (2014). A Toolbox for Representational Similarity Analysis. PLoS Computational Biology, 10(4), e1003553. https://doi.org/10.1371/journal.pcbi.1003553

Noble, K. G., Houston, S. M., Brito, N. H., Bartsch, H., Kan, E., Kuperman, J. M., … Sowell, E. R. (2015). Family income, parental education and brain structure in children and adolescents. Nature Neuroscience, 18, 773–778.

Oedekoven, C. S. H., Keidel, J. L., Berens, S. C., & Bird, C. M. (2017). Reinstatement of memory representations for lifelike events over the course of a week. Scientific Reports. https://doi.org/10.1038/s41598-017-13938-4

Owen, A. M., McMillan, K. M., Laird, A. R., & Bullmore, E. (2005). N-back working memory paradigm: A meta-analysis of normative functional neuroimaging studies. Human Brain Mapping, 25(1), 46–59. https://doi.org/10.1002/hbm.20131

Pallier, C. (2002). Computing discriminability and bias with the R software. Retrieved from http://citeseerx.ist.psu.edu/viewdoc/download?doi=10.1.1.218.7769&rep=rep1&type=pdf

Pinel, P., Lalanne, C., Bourgeron, T., Fauchereau, F., Poupon, C., Artiges, E., … Dehaene, S. (2015). Genetic and environmental influences on the visual word form and fusiform face areas. Cerebral Cortex, 25(9), 2478–2493. https://doi.org/10.1093/cercor/bhu048

Plomin, R., & Deary, I. J. (2015). Genetics and intelligence differences: five special findings. Molecular Psychiatry, 20, 98–108.

Plomin, R., Willerman, L., & Loehlin, J. C. (1976). Resemblance in appearance and the equal environments assumption in twin studies of personality traits. Behavior Genetics, 6(1), 43–52. https://doi.org/10.1007/BF01065677

Polderman, T. J. C., Benyamin, B., de Leeuw, C. A., Sullivan, P. F., van Bochoven, A., Visscher, P. M., & Posthuma, D. (2015). Meta-analysis of the heritability of human traits based on fifty years of twin studies. Nature Genetics, 47(7), 702–709. https://doi.org/10.1038/ng.3285

Polk, T. A., Park, J., Smith, M. R., & Park, D. C. (2007). Nature versus nurture in ventral visual cortex: a functional magnetic resonance imaging study of twins. The Journal of Neuroscience : The Official Journal of the Society for Neuroscience, 27(51), 13921–13925. https://doi.org/10.1523/JNEUROSCI.4001-07.2007

R Development Core Team. (2015). R: A language and environment for statistical computing. Vienna, Austria: R Foundation for Statistical Computing.

Reverberi, C., Görgen, K., & Haynes, J.-D. (2012). Compositionality of Rule Representations in Human Prefrontal Cortex. Cerebral Cortex, 22(6), 1237–1246. https://doi.org/10.1093/cercor/bhr200

Riggall, A. C., & Postle, B. R. (2012). The Relationship between Working Memory Storage and Elevated Activity as Measured with Functional Magnetic Resonance Imaging. The Journal of Neuroscience, 32(38), 12990–12998. https://doi.org/10.1523/jneurosci.1892-12.2012

Romesburg, C. (2004). Cluster Analysis for Researchers. Lulu.com.

Rottschy, C., Langner, R., Dogan, I., Reetz, K., Laird, A. R., Schulz, J. B., … Eickhoff, S. B. (2012). Modelling neural correlates of working memory: A coordinate-based meta-analysis. NeuroImage, 60(1), 830–846. https://doi.org/https://doi.org/10.1016/j.neuroimage.2011.11.050

Satterthwaite, T. D., Xia, C. H., & Bassett, D. S. (2018). Personalized Neuroscience: Common and Individual-Specific Features in Functional Brain Networks. Neuron. https://doi.org/10.1016/j.neuron.2018.04.007

Serences, J. T., Ester, E. F., Vogel, E. K., & Awh, E. (2009). Stimulus-Specific Delay Activity in Human Primary Visual Cortex. Psychological Science, 20(2), 207–214. https://doi.org/10.1111/j.1467-9280.2009.02276.x

Smith, S. M., Jenkinson, M., Woolrich, M. W., Beckmann, C. F., Behrens, T. E. J., Johansen-Berg, H., … Matthews, P. M. (2004). Advances in functional and structural MR image analysis and implementation as FSL. NeuroImage, 23 Suppl 1, S208–19. https://doi.org/10.1016/j.neuroimage.2004.07.051

Tavor, I., Parker Jones, O., Mars, R. B., Smith, S. M., Behrens, T. E., & Jbabdi, S. (2016). Task-free MRI predicts individual differences in brain activity during task performance. Science. https://doi.org/10.1126/science.aad8127

Van Essen, D. C., Ugurbil, K., Auerbach, E., Barch, D., Behrens, T. E. J., Bucholz, R., … Yacoub, E. (2012). The Human Connectome Project: A data acquisition perspective. Neuroimage, 62(4), 2222–2231. https://doi.org/http://dx.doi.org/10.1016/j.neuroimage.2012.02.018

Verhulst, B., & Hatemi, P. K. (2013). Gene-Environment Interplay in Twin Models. Political Analysis, 21(3), 368–389. https://doi.org/DOI:10.1093/pan/mpt005

Vincent, J. L., Kahn, I., Snyder, A. Z., Raichle, M. E., & Buckner, R. L. (2008). Evidence for a Frontoparietal Control System Revealed by Intrinsic Functional Connectivity. Journal of Neurophysiology, 100(6), 3328–3342. https://doi.org/10.1152/jn.90355.2008

Waskom, M. L., Kumaran, D., Gordon, A. M., Rissman, J., & Wagner, A. D. (2014). Frontoparietal Representations of Task Context Support the Flexible Control of Goal-Directed Cognition. The Journal of Neuroscience, 34(32), 10743–10755. https://doi.org/10.1523/jneurosci.5282-13.2014

Wilcox, R. R. (2017). Introduction to robust estimation and hypothesis testing. (4th ed.). Elsevier Academic Press.

Woolgar, A., Hampshire, A., Thompson, R., & Duncan, J. (2011). Adaptive Coding of Task-Relevant Information in Human Frontoparietal Cortex. The Journal of Neuroscience, 31(41), 14592–14599. https://doi.org/10.1523/jneurosci.2616-11.2011

Woolgar, A., Thompson, R., Bor, D., & Duncan, J. (2011). Multi-voxel coding of stimuli, rules, and responses in human frontoparietal cortex. Neuroimage, 56(2), 744–752. https://doi.org/http://dx.doi.org/10.1016/j.neuroimage.2010.04.035

Xue, G., Dong, Q., Chen, C., Lu, Z., Mumford, J. A., & Poldrack, R. A. (2010). Greater Neural Pattern Similarity Across Repetitions Is Associated with Better Memory. Science, 330(6000), 97–101. https://doi.org/10.1126/science.1193125

Zhang, J., Kriegeskorte, N., Carlin, J. D., & Rowe, J. B. (2013). Choosing the Rules: Distinct and Overlapping Frontoparietal Representations of Task Rules for Perceptual Decisions. The Journal of Neuroscience, 33(29), 11852–11862. https://doi.org/10.1523/jneurosci.5193-12.2013

